# A Common PD-Risk *GBA1* Variant Disrupts LIMP2 Interaction, Impairs Glucocerebrosidase Function, and Drives Lysosomal and Mitochondrial Dysfunction

**DOI:** 10.1101/2025.08.28.672891

**Authors:** Oliver B. Davis, Jennifer E. Kung, Sonnet S. Davis, Rajarshi Ghosh, Shan V. Andrews, Neal S. Gould, Jillian H. Kluss, Elliot Thomsen, Maayan Agam, John P. Coan, Michael T. Maloney, Ann H. Nguyen, Hoang N. Nguyen, Nicholas E. Propson, Edwin I. Lozano, Tianao Yuan, Kaitlin Xa, R. Andres Parra Sperberg, Shourya Jain, Roger Lawrence, Julie C. Ullman, Srijana Balasundar, H. Paul Benton, Maja Petkovic, Ahlam N. Qerqez, Xiang Wang, Sha Zhu, Gilbert Di Paolo, Mihalis S. Kariolis, Cathal S. Mahon, Annie Arguello, David J. Vocadlo, Mark R. Cookson, Jung H. Suh, Lionel Rougé, Anastasia G. Henry

**Affiliations:** Denali Therapeutics Inc., South San Francisco, CA, United States; National Institute on Aging, National Institutes of Health, Bethesda, MD, United States; Department of Chemistry, Simon Fraser University, Burnaby, BC, Canada; Tenvie Therapeutics Inc., South San Francisco, CA, United States

## Abstract

Variants in *GBA1* cause Gaucher disease (GD), a lysosomal storage disorder, and represent the most common genetic risk factor for Parkinson’s disease (PD). While some *GBA1* variants are associated with both GD and PD, several coding mutations, including E326K, specifically confer risk for developing PD. It is established that GD-linked variants in β-glucocerebrosidase (GCase), the enzyme encoded by *GBA1*, are loss-of-function, but it remains unclear whether variants solely associated with PD similarly reduce GCase activity. The mechanisms by which some of these variants impact GCase activity and PD-associated pathways, including lysosomal and mitochondrial function, are also poorly defined. Here, we show that the PD-linked E326K variant significantly reduces lysosomal GCase activity by impairing its delivery to lysosomes via altered interactions with its receptor, LIMP2. Biophysical and structural characterization of this variant, both alone and in complex with LIMP2, reveals a dimeric organization that appears to result from the loss of a key salt bridge between E326 and R329. Restoration of this salt bridge through the introduction of a negatively charged side chain at position 329 promotes monomeric organization and interaction with LIMP2 in cells. *GBA1*-p.E326K cell models show greater deficits in PD-linked pathways compared to more severe loss of GCase function, including secondary lysosomal lipid storage and mitochondrial dysfunction. We confirm the E326K variant impacts GCase pathway activity in relevant CNS cell types, including iPSC-derived microglia, and in biofluids from heterozygous *GBA1-*p.E326K variant carriers. Together, our data provide key insights into the nature of GCase dysfunction in *GBA1*-PD and can inform the development of GCase-targeted therapeutic strategies to treat PD.

## Introduction

*GBA1* encodes glucocerebrosidase (GCase), a lysosomal enzyme that catalyzes the hydrolysis of its substrates glucosylceramide (GlcCer) and glucosylsphingosine (GlcSph)^1^. Homozygous or biallelic loss-of-function (LoF) mutations in *GBA1* cause the lysosomal storage disorder (LSD) Gaucher Disease (GD), which is characterized by accumulation of GlcCer and GlcSph and subsequent lysosomal dysfunction in affected cells and tissues of GD patients.^2^ Mechanistically, GD-associated *GBA1* coding variants are thought to cause GCase LoF by producing mutant and/or misfolded enzymes with reduced lysosomal GCase activity.^3–5^ While significant phenotypic heterogeneity exists in the clinical presentation of GD, the degree to which a given mutation reduces lysosomal GCase activity is correlated to some extent with the severity of disease manifestation and/or neurological involvement.^6–8^ Mutations in the *GBA1* gene have also been identified as the most common genetic risk factor for Parkinson’s Disease (PD), with estimated frequencies ranging from 5-20% of PD patients carrying at least one *GBA1* variant allele.^9–11^ Intriguingly, several variants in *GBA1* have been identified that are associated with increased risk for PD but do not appear to cause GD.^9,12,13^ The existence of these PD-specific *GBA1* variants raises the question of whether common or distinct mechanisms for GCase dysfunction underlie GD and *GBA1*-dependent PD pathologies. One of the most common coding variants in *GBA1* associated with PD risk is the E326K variant (also annotated as E365K using the naming convention that includes the 39 amino acid leader sequence), which is highly enriched in certain PD patient cohorts, with a general population frequency of approximately 1.2% across ancestral populations and reported odds ratios for PD between 1.57 and 5.50.^9,12,14,15^ While GD-associated variants have been shown to lead to a severe impairment in GCase activity^2^, the consequences of E326K remain poorly defined as conflicting results have been reported on if and how this variant impacts lysosomal GCase activity.^16–20^

Reduced GCase activity has been mechanistically linked to various forms of lysosomal dysfunction^21–26^, and emerging evidence from human genetics studies and analysis of PD patient samples has broadly implicated lysosomal dysfunction as a key driver of disease.^27,28^ In addition to *GBA1*, various genetic association studies have nominated variants in genes involved in endolysosomal and/or autophagic function as PD-risk factors, including *LRRK2*, *TMEM175*, and *ATP13A2*.^29–34^ Furthermore, an increased burden of rare LSD variants has been characterized in PD patient populations, and activity of multiple lysosomal hydrolases, including GCase itself, have been shown to be reduced in post-mortem analysis of PD patient brains, adding credence to the notion that endolysosomal dysfunction contributes to PD pathogenesis.^35–37^ The consequences of PD-specific *GBA1* variants, such as E326K, on lysosomal function and cellular homeostasis more broadly are poorly understood. A deeper understanding of how common PD-associated variants like E326K impact lysosomal function is paramount as there are currently no approved disease-modifying therapies for PD. Moreover, profiling dysfunction downstream of *GBA1* is essential to help identify pathway biomarkers that can enable the clinical development of GCase-targeted therapeutic approaches.

Here, we investigated the impact of the E326K variant using a comprehensive approach that employed analysis of recombinant enzyme, preclinical cell and animal models, and human samples from *GBA1*-p.E326K variant carriers to delineate how this variant affects GCase structure and lysosomal activity, as well as PD-relevant aspects of lysosomal and mitochondrial function. We demonstrate that the E326K variant causes GCase LoF that stems from reduced transport of this variant enzyme to the lysosome, and we show that this impairment results from reduced interactions between mutant GCase and its receptor, LIMP2. Biophysical and structural studies indicate that the E326K GCase variant, unlike the WT enzyme, behaves as a constitutive dimer in solution. We establish ^38–40,41^ that dimerization of E326K results from the loss of a key salt bridge between the side chains of E326 and R329, and we show that introduction of a compensatory charge pair in the E326K variant can restore the monomeric state of the enzyme as well as its ability to interact with LIMP2 in cells. Using lipidomic and proteomic profiling of purified lysosomes, we found that the overall effect of the E326K variant on lysosomal GCase enzyme levels and substrate accumulation is mild in comparison to the L444P variant, a well-characterized severe LoF mutation that is associated with neuronopathic GD. Unexpectedly, our results reveal that the E326K variant leads to greater secondary lipid storage and mitochondrial dysfunction than the L444P variant despite its milder GCase LoF phenotype. We demonstrate that these findings translate to CNS-relevant models and cell types, including E326K knock-in mouse brains and human iPSC-derived microglia. Additionally, we observed reduced GCase activity and accumulation of GCase substrates in biofluids from human subjects that are heterozygous carriers of the *GBA1*-p.E326K variant. Our results identify candidate lipid biomarkers of secondary pathway dysfunction in samples from E326K variant carriers, and we demonstrate that recombinant GCase is sufficient to reduce accumulation of these lipid biomarkers in E326K cellular models. Collectively, these results improve our mechanistic understanding of how reduced GCase activity can contribute to key aspects of PD pathogenesis and highlight the therapeutic potential of approaches aimed at restoring GCase levels for the treatment of PD.

## Results

### GBA1-p.E326K causes GCase loss-of-function via reduced lysosomal trafficking

To evaluate the impact of endogenous expression of the E326K variant on GCase activity and understand its effects in comparison to GD-linked variants in *GBA1*, we generated a suite of knock-in (KI) HEK293T cell lines using CRISPR/Cas9 that express the E326K variant or the GD-associated severe LoF mutation L444P. *GBA1* knock-out (KO) cells were also made to control for total loss of GCase activity. Using a recently developed substrate-mimetic probe to measure endolysosomal GCase activity (LysoFQ-GBA),^38^ we observed reduced GCase activity in all variant cells compared to WT cells and confirmed lack of signal in the *GBA1* KO cells.

E326K KI clones retained higher levels of GCase activity compared to L444P KI clones, with an average of 51.8% of WT activity as opposed to the 3.2% retained activity in L444P KI cell lines (Fig. 1A, B).

**Figure 1.**
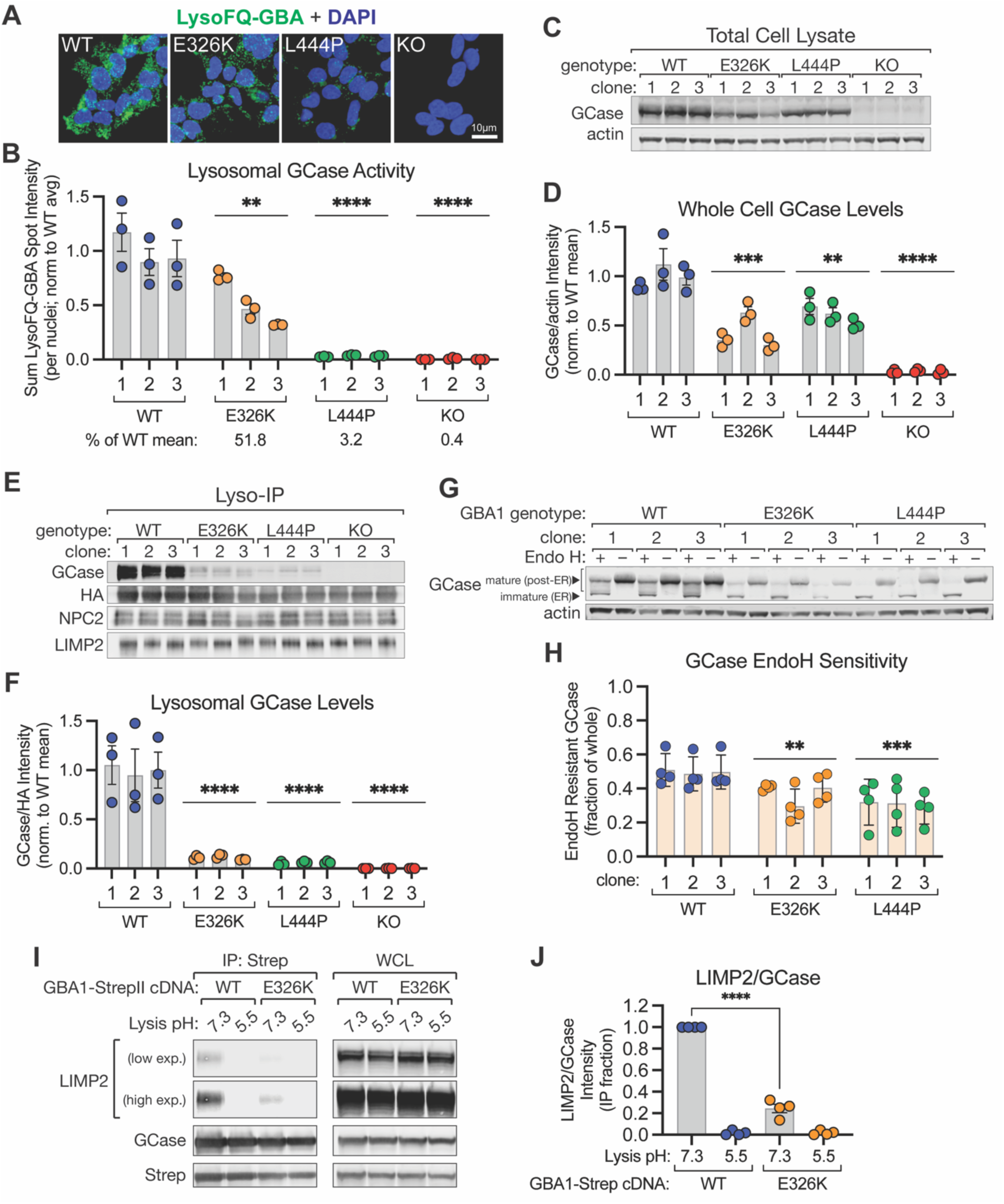
*GBA1*-p.E326K causes a loss of GCase activity in cells through impaired lysosomal targeting. **A** Representative micrographs of data quantified in **B** from WT, *GBA1*-p.E326K KI, *GBA1*-p.L444P KI, and *GBA1* KO cells depicting LysoFQ-GBA probe fluorescence (green). Nuclei stained with DAPI (blue); scale bar is 10 µm. **B** WT, *GBA1*-p.E326K KI, *GBA1*-p.L444P KI, and *GBA1* KO HEK293T cells were stained with LysoFQ-GBA probe to measure lysosomal GCase activity, and the mean GCase activity for each clone is quantified as the sum LysoFQ-GBA spot intensity per nuclei, normalized to the mean of all WT cells. n = 3 independent experiments. **C** Representative immunoblot of whole cell lysates from HEK293T cells of indicated genotype assessed with antibodies against GCase and actin as a loading control. **D** Quantification of total cellular GCase enzyme levels, measured via immunoblot & normalized to the levels of actin and the mean signal from WT cells. n = 3 independent experiments. **E** Representative immunoblot of Lyso-IP samples from HEK293T cells (of indicated genotype) expressing TMEM192-HA^x3^ tag. Immunoblots were probed using antibodies against GCase, HA, NPC2, and LIMP2. **F** Quantification of lysosomal GCase levels (normalized per sample to HA band intensity), measured via immunoblotting of Lyso-IP fraction and normalized to the mean of all WT samples. n = 4 independent experiments. **G** Representative immunoblot of total cell lysates from HEK293T cells of indicated genotype, treated ± Endoglycosidase H (EndoH), and probed using antibodies against GCase and actin as a loading control. **H** Quantification of the EndoH-resistant GCase band intensity normalized to the total GCase band intensity, from EndoH-treated cell lysates from HEK293T cells of indicated genotype. n = 4 independent experiments. **I** Representative immuoblots from Strep-Tactin pulldowns performed on lysates from *GBA1* KO HEK293T cells transfected with StrepII-tagged *GBA1* cDNA for WT *GBA1* and *GBA1*-p.E326K. Immunoblots were probed using antibodies against LIMP2, GCase and Strep. **J** Quantification of LIMP2:GCase band intensity ratios, normalized to the WT GBA1/pH 7.3 lysis condition within each replicate, from pulldown fraction of experiment shown in I. n = 4 independent experiments. All bar graphs depict mean ± SEM with individual points representative of data from a single experimental replicate. Unless otherwise noted, all statistics were performed on genotype-level comparisons using one-way ANOVA with Dunnett’s multiple comparison test, where ** = p < 0.0021, *** = p < 0.0002, **** = p < 0.0001.

To better define the mechanism underlying the loss of GCase activity in E326K KI cells, we assessed GCase levels in both whole cell lysates (Fig. 1C, D) and in lysosomes isolated from these cells via rapid immunopurification using the established Lyso-IP method^39^ (Fig. 1E, F & S1A, B). GCase levels were reduced at the whole cell level in both E326K and L444P KI cells compared to WT cells, with an even greater relative reduction of enzyme levels observed in isolated lysosomes from cells expressing these variants, suggesting that the E326K variant may negatively impact the delivery of GCase to lysosomes. As the L444P variant has been previously shown to misfold in the ER and be prematurely degraded before being delivered to the lysosome^5^, we decided to investigate the trafficking of the E326K variant to determine if a similar mechanism may be involved in reducing lysosomal enzyme levels. We employed the classical endoglycosidase H (endo H) sensitivity assay to examine whether the E326K variant was more sensitive to endo H treatment, indicative of defects in trafficking out of the ER and early Golgi complex.^40–43^ Analysis of the electrophoretic mobility of endogenous GCase revealed higher endo H sensitivity of the E326K variant compared to WT enzyme, indicating reduced glycan complexity, and suggesting that the E326K variant has impaired trafficking out of the biosynthetic pathway (Fig. 1G, H). Higher endo H sensitivity was observed with the endogenous L444P enzyme compared to E326K, correlating well with our results showing a stronger reduction in lysosomal enzyme levels and activity. These results implicate defective lysosomal trafficking as a common underlying mechanism of GCase LoF.

GCase requires interactions with its receptor, LIMP2, to be efficiently trafficked to the lysosome,^44,45^ and we hypothesized that the E326K variant may be inefficiently delivered to the endolysosomal compartment due to altered interactions with LIMP2. We assessed the interaction between the E326K variant enzyme and LIMP2 via co-immunoprecipitation and found that the E326K variant interacts less efficiently with LIMP2 at neutral pH than the WT enzyme does upon overexpression in *GBA1* KO cells (Fig. 1I, J and S1C). Under acidic conditions mimicking lysosomal pH, both WT and E326K variant GCase exhibit lack of binding to LIMP2, suggesting that cellular interaction between LIMP2 and GCase appears to be pH sensitive. While this observation is consistent with previous reports^46–48^, recent studies showed that LIMP2 can still bind and activate GCase under lysosomal conditions^45,49^. Together, these results suggest that the E326K variant reduces lysosomal GCase activity by disrupting its interaction with LIMP2 to ultimately impair GCase delivery to lysosomes.

### GCase E326K does not impact enzymatic activity in vitro

To better understand how the E326K variant leads to impaired interaction with LIMP2 in cells, we recombinantly expressed and purified the mutant protein and compared its biophysical properties and interaction with recombinant LIMP2 to that of recombinant WT GCase.

Expression yields for both WT and E326K variant GCase were comparable, and both proteins were purified to homogeneity. To determine potential effects of the E326K variant on the enzyme’s intrinsic activity, we profiled the enzymatic activity of WT and E326K variant GCase proteins. Both proteins exhibited nearly identical enzymatic activity using an established 4-MU-β-Glc based-assay^50^ (Fig. 2A, B, Fig. S2A). We also employed a recently developed liposome-based assay^51^ designed to monitor GCase activity against its native substrate GlcCer embedded in a membrane-like environment. In this more physiologically-relevant setting, the E326K variant also exhibited a catalytic profile that was comparable to that of WT GCase and imiglucerase – a recombinant GCase that is used as an enzyme replacement therapy for Gaucher Disease (Fig 2C). Changes to liposomal lipid composition did not differentially affect the catalytic activity of any of the recombinant enzymes tested, as inclusion of 30% bis(monoacylglycero)phosphate (BMP, an anionic phospholipid that can stimulate GCase activity to promote GSL hydrolysis^52–55^) equally increased the specific activity of both WT and E326K variant GCase (Fig. 2D). Collectively, these results suggest that the reduced lysosomal GCase activity observed in our *GBA1*-p.E326K KI cell model likely stems from impaired lysosomal delivery of GCase rather than defects in the intrinsic activity or substrate recognition of the variant enzyme.

**Figure 2.**
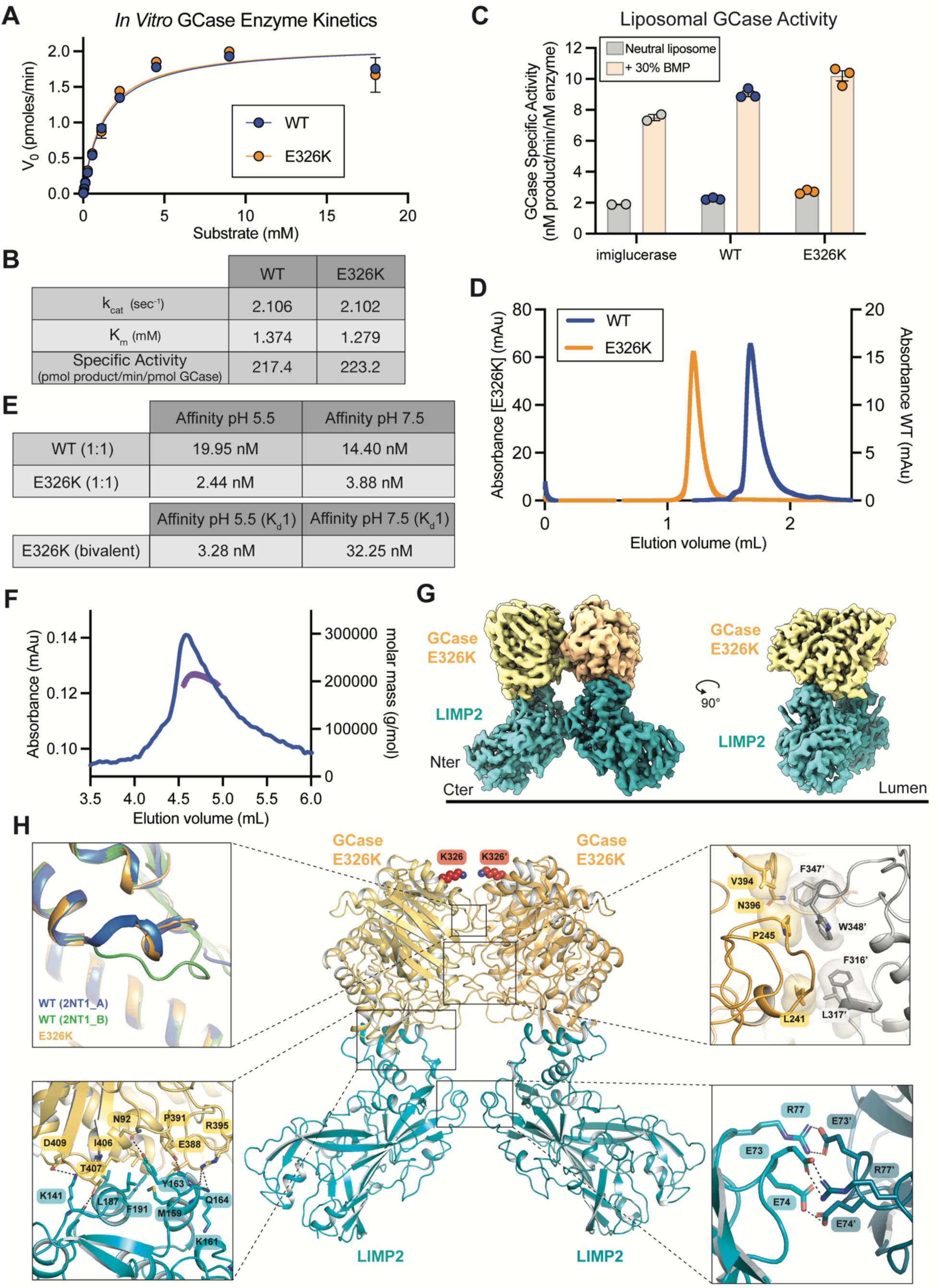
Biochemical and structural characterization of the E326K variant and in vitro interaction with LIMP2. **A** Enzymatic activity of purified recombinant GCase (WT or E326K) against the substrate 4-methylumbelliferyl b-glucopyranoside (4-MUG), at an enzyme concentration of 2 nM. Curves depict data fits to Michaelis-Menten kinetics model (top). **B** Selected enzymatic parameters, derived from Michaelis-Menten kinetics shown in **B**. **C** Enzymatic activity of purified recombinant GCase (WT or E326K) against GlcCer in liposomes prepared with either neutral lipids (gray bars) or with 30% BMP (orange bars). Imiglucerase is colored in grey, recombinant WT GCase in blue and E326K in orange. n = 3 independent experiments. All bar graphs depict mean ± SEM with individual points representative of data from a single experimental replicate. **D** Overlay of size exclusion chromatography profiles of purified WT (blue) and E326K (orange) GCase. Samples were run over a Superdex S200 3.2/100 pre-equilibrated in 20mM sodium acetate pH 5.5 and 150mM NaCl and absorbance at 280nm was recorded. Early elution profile for E326K suggests that protein behaves as a multimer compared to WT GCase. Void volume is expected around 0.9mL confirming E326K is not a soluble aggregate. **E** Binding affinities measured by SPR for the LIMP2-GCase interaction at both neutral and acidic pHs, with LIMP2 lumenal domain immobilized on the chip. Values correspond to the average of 2 independent replicates, using either a 1:1 (top table) or biphasic (bottom table) binding model. **F** SEC-MALS analysis of the GCase E326K / LIMP2 complex is consistent with a 2:2 dimeric organization (observed molecular weight ∼ 200 kDa, expected MW for GCase ∼50kDa and LIMP2 extracellular domain ∼45kDa, not accounting for glycosylation). Absorbance trace is shown in blue, and apparent molecular weight is shown in purple. **G** CryoEM reconstruction of the E326K variant & LIMP2 lumenal domain complex at a resolution of 3.3 Å. Two orthogonal side views of an isosurface rendering of the complex are shown. LIMP2 N- and C-termini are indicated relative to the lysosomal membrane. **H** Cartoon representation of the E326K variant & LIMP2 complex structure (center). K326 in both E326K variant protomers is shown in red spheres. Overlay of Loop 1 in E326K variant & LIMP2 complex structure with helical (blue) and extended (green) conformations of Loop 1 observed in WT GCase at neutral pH (PDB: 2NT1, chains A and B) and (top left). Close-up view of E326K dimer interface showing that the interface consists primarily of hydrophobic interactions (upper right). Close-up view of interactions forming the E326K variant interface with LIMP2 (bottom left). Close-up view of the electrostatic interactions at the LIMP2 dimer interface (bottom right).

### GCase E326K forms a constitutive dimer and a 2:2 complex with LIMP2

Next, we further characterized the biophysical and biochemical properties of the E326K GCase variant to identify features that might underlie its impaired trafficking and reduced lysosomal activity. Interestingly, we noticed during the purification process that its size exclusion chromatography (SEC) profile suggested the E326K variant behaves as a dimer at both acidic and neutral pH, whereas WT GCase is primarily monomeric (Fig. 2D, S2B). Dynamic light scattering (DLS) analysis further confirmed that the E326K variant behaves as a constitutive dimer in solution, exhibiting a radius approximately twice that of WT GCase (Fig S2C, D).

Differential scanning fluorimetry (DSF) analysis also showed that the melting temperature (T_m_) for the E326K dimer was higher than that of the WT protein at both neutral and acidic pH, suggesting that the multimeric organization of the variant led to increased stability (Fig. S2C, D). These results indicate that the observed E326K dimer is not pH sensitive and is stable at both neutral and acidic pH. To investigate the impact of GCase E326K dimerization on LIMP2 binding, we purified recombinant human LIMP2 lumenal domain (residues 36-430) and assessed its ability to interact with WT and E326K variant GCase through surface plasmon resonance (SPR) at both neutral and acidic pH using a biotinylated LIMP2 construct (Fig. 2E and Fig. S2E, F). We first tested binding between WT GCase and LIMP2 and observed a modest decrease between binding at pH 5.5 and pH 7.5 (K_d_ = 19.95 nM and 14.40 nM, respectively), using a 1:1 binding model. While our finding is contrary to our earlier observation that this interaction appears to be weaker at acidic pH in a cellular environment, the largely unchanged affinity of the GCase/LIMP2 complex at acidic pH in vitro could be due to unknown factors that play a role in modulating this interaction in a physiological setting (e.g. at the endolysosomal membrane).

While a 1:1 binding model determined that the binding affinity of E326K to LIMP2 was comparable to that of WT GCase (K_d_ = 2.44 nM at pH 5.5 & 3.89 nM at pH 7.5), this model does not account for E326K dimers displaying two potential LIMP2 binding sites. Therefore, we instead used a bivalent binding model to more accurately capture the sequential binding and possible avidity effects that influence the kinetics of this interaction. Using this bivalent model, the first site dissociation constant (K_d_1) of the E326K-LIMP2 interaction was ∼2x weaker than the monovalent 1:1 dissociation constant (K_d_) of the WT-LIMP2 interaction at neutral pH (Fig. 2E, Fig. S2E, F).

We sought to further investigate the molecular basis underlying how the E326K mutation and its propensity to dimerize leads to altered interactions with LIMP2 by determining structures of complexes between LIMP2 and WT or E326K variant GCase through cryo-electron microscopy (cryoEM). Consistent with the affinities measured via SPR, we were able to form stable complexes of LIMP2 bound to E326K variant GCase by SEC (Fig. S2F). Using SEC coupled with Multi-Angle Light Scattering (SEC-MALS), we confirmed that the E326K variant forms a 2:2 complex with LIMP2 (Fig. 2F). For the GCase E326K-LIMP2 complex (Fig. S2F), we obtained a 3D reconstruction with an overall resolution of 3.3 Å (Fig. S3A-E), revealing a symmetric 2:2 heterotetramer (Fig. 2G, Table S1), in which both GCase E326K and LIMP2 participate in homotypic, as well as heterotypic, interactions. The two GCase E326K molecules form a symmetric homodimer with a buried surface area of 519 Å^2^ (Fig. 2G, H). These findings indicate that this GCase mutant behaves as a constitutive dimer, revealing a potential new LoF mechanism and highlighting the importance of the monomer–dimer equilibrium in GCase function. GCase dimers have previously been observed in several crystal structures in both acidic and neutral conditions, as well as in the presence or absence of small molecule activators or inhibitors^56–58^. Consistent with prior GCase structures^59^, the dimer observed in the E326K variant-LIMP2 structure positions the active sites of both monomers facing each other such that the dimer interface consists primarily of hydrophobic contacts mediated by loops surrounding the active sites (Fig. 2H). These loops have been proposed to be involved in ceramide moiety binding, and Loop 1 (residues 311-319) is thought to rearrange from an extended to a helical conformation in the active state of the enzyme^56^. Notably, Loop 1 in the E326K variant adopts a helical conformation (Fig. 2H), consistent with this variant maintaining enzymatic activity.

Variations in the orientation of the GCase monomers relative to each other have been observed in prior GCase dimer structures, and in our E326K variant structure we observe a distinct arrangement of the monomers that differs from previous structures of WT GCase or the N370S variant (Fig. S4A). ^61^Interestingly, the E326K mutation itself is located near the dimer interface but does not directly form interactions with the other protomer (Fig. 2H). Instead, this residue is part of an α-helix C-terminal to Loop 1, which is directly involved in the dimer interface.

The conformation of LIMP2 in the complex resembles that observed in published crystal structures^60–62^. Strikingly, however, dimerization of GCase E326K positions the two LIMP2 molecules such that they form homotypic interactions, resulting in a small interface (BSA = 278 Å^2^) made up of electrostatic interactions between E73 in one protomer and R77 of the other, as well as hydrogen bonds between E74 of one protomer and the main chain of T70 in the other (Fig. 2H). Similar interfaces are present as crystal contacts in several crystal structures of LIMP2, consistent with this region having some propensity to mediate self-association (Fig. S4B).^62^ Due to the small size of this interface, however, it seems unlikely that LIMP2 would homodimerize through this site in the absence of interaction with the GCase E326K dimer.

Alignment of the GCase E326K-LIMP2 structure with the recently published cryoEM structure of a nanobody-bound GCase WT-LIMP2 complex reveals small conformational differences in both proteins (Fig. S3F)^45^. However, the interface between GCase E326K and LIMP2 closely resembles that observed in the GCase WT-LIMP2 complex, with both interfaces consisting mostly of hydrophobic interactions and a few polar interactions (Fig. 2H). Alignment of two copies of the GCase WT-LIMP2 complex with the GCase E326K-LIMP2 heterotetramer suggests that the conformations of GCase WT and LIMP2 are compatible with dimerization (Fig. S3G). While we were unable to obtain a high-resolution reconstruction of the GCase WT-LIMP2 complex due to strong preferred orientation, 2D classes and low-resolution reconstructions confirmed formation of a 1:1 heterodimer (Fig. S3H) that is consistent with the recently published structure of this complex^45^. The absence of higher order complexes of WT GCase and LIMP2 in both our cryoEM and biophysical studies further underscores that formation of the 2:2 GCase E326K-LIMP2 heterotetramer is driven by dimerization of the E326K variant.

Furthermore, the lack of major differences in the interaction interfaces of GCase WT and E326K with LIMP2 suggests that this change in stoichiometry of the complex due to E326K dimerization might underlie the functional defects observed for this variant.

### A key salt bridge between E326 and R329 stabilizes GCase in a monomeric state and restores interaction with LIMP2 in cells

Based on our biophysical and structural characterization of the GCase E326K-LIMP2 interaction, we hypothesized that the dimeric nature of the E326K variant is a key determinant contributing to its impaired lysosomal trafficking. To further understand the molecular basis for this dimerization, we solved a 3.1 Å crystal structure of the E326K variant (Fig. 3A, Table S2).

**Figure 3.**
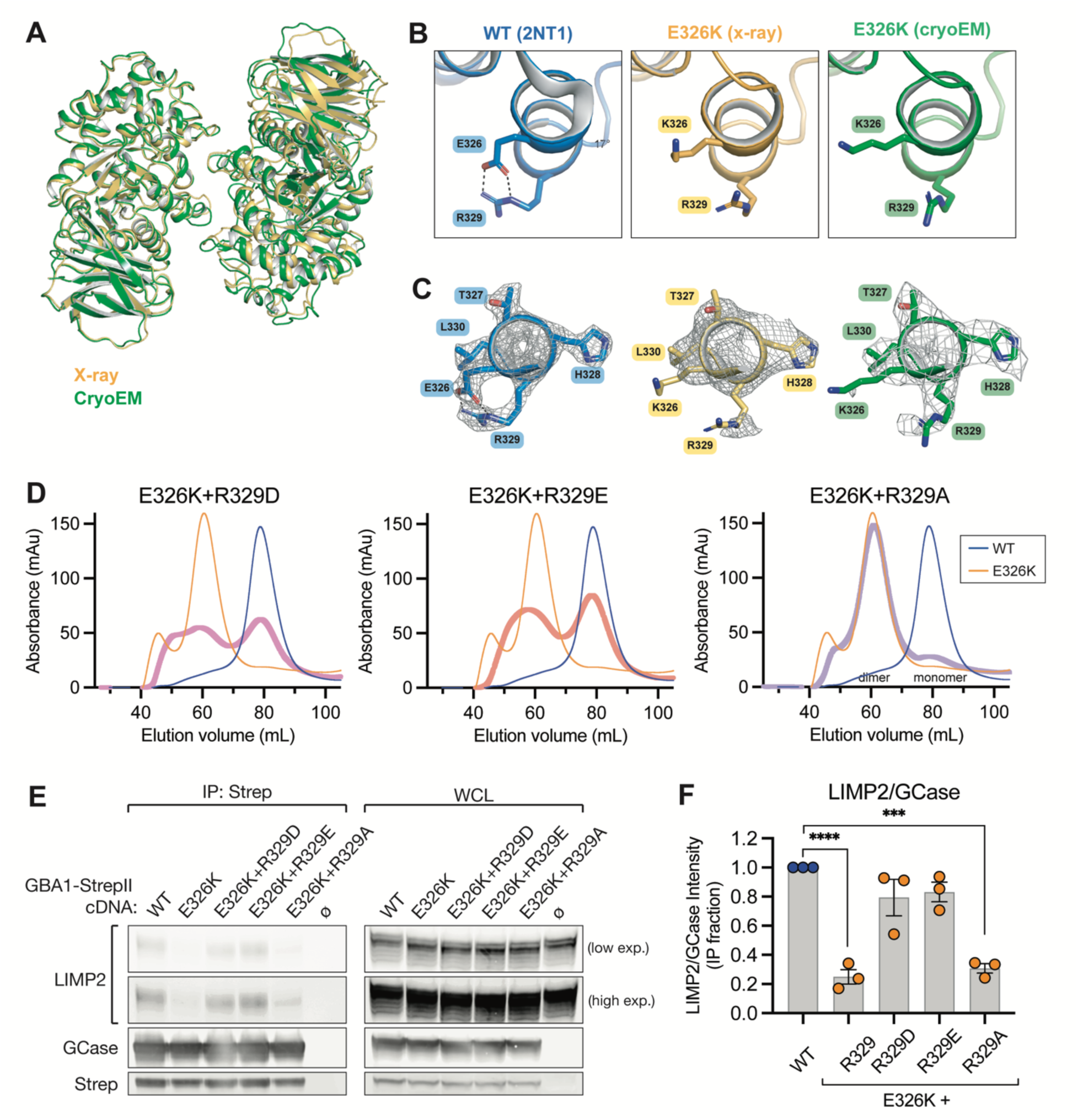
A salt bridge between residues 326 and 329 regulates GCase self-association and binding to LIMP2 in cells. **A** Crystal structure of the GCase E326K variant dimer at 3.1 Å. Overlay of the E326K variant dimer observed in the crystal structure of E326K alone (gold) and the cryoEM structure of the E326K variant & LIMP2 complex (green). **B** Close-up view of the interactions of residue 326 in GCase structures. In WT GCase (PDB:2NT1, blue) the carboxylic side chain of Glu326 engages in a salt bridge interaction with guanidium group of R329. This electrostatic interaction is lost in both the E326K variant crystal structure (gold) and the cryoEM structure of the E326K variant & LIMP2 complex (green). **C** Corresponding electron density (mesh representation) for panel B (contoured at 1σ for the WT and E326K crystal structures, and 8σ for the E326K/LIMP2 cryoEM structure), showing that the E326 and R329 sidechains are both well-ordered and well-resolved in WT GCase, while the K326 and R329 sidechains are more disordered in the E326K variant. **D** Size exclusion chromatography profiles of purified E326K+R329A (purple), E326K+R329E (light red), and E326K+R329A (violet) recombinant enzyme. Elution profiles for WT GCase and E326K variant are shown in blue and orange, respectively. Elution profiles of monomeric and dimeric proteins are indicated with text. **E** Representative immunoblot from Strep-Tactin pulldowns performed on lysates from *GBA1* KO HEK293T cells transfected with StrepII-tagged GBA1 cDNA for WT *GBA1*, *GBA1*-p.E326K, and E326K+R329D/E/A variants. Immunoblots were probed using antibodies against LIMP2, GCase and Strep. **F** Quantification of LIMP2:GCase band intensity ratios, normalized to the WT *GBA1* transfected condition within each replicate, from pulldown fraction of experiment shown in **E**. n = 3 independent experiments, statistics performed using one-way ANOVA with Dunnett’s multiple comparison test to WT GCase, where *** = p < 0.0002, **** = p < 0.0001. All bar graphs depict mean ± SEM with individual points representative of data from a single experimental replicate.

The asymmetric unit contained four copies of GCase E326K, forming a dimer of dimers (Fig. S5A), with both dimers having a buried surface area (BSA) of about 445 Å^2^. The predominant dimer adopts a conformation similar to that observed in the GCase E326K-LIMP2 cryoEM structure where both active sites are facing each other. Overall, the dimers observed in both the cryoEM and X-ray structures align closely with a RMSD of 2.5 Å (Fig. 3A). The slight differences observed may result from crystal packing, which could affect the orientation of the protomers relative to each other. Since the E326K mutation is distal from the dimer interface, we examined the local environment surrounding K326 more closely to identify possible structural changes that might contribute to the altered oligomeric state of the E326K variant. Comparison with structures of WT GCase revealed that the negatively-charged carboxylic acid sidechain of E326 typically forms an electrostatic interaction with the positively-charged guanidium group of R329 and that this interaction is absent in both our crystal and cryoEM structures of the E326K variant (Fig. 3B, C). Intramolecular salt bridges within α-helices have been shown to speed up protein folding rates and more generally to stabilize final protein conformations.^63^ Here, the enthalpic cost related to the loss of this salt bridge might result in an increased intrinsic instability of the mutant, promoting GCase dimerization through subtle allosteric changes at the dimer interface to compensate for this greater instability.

To directly test whether loss of the salt bridge between positions 326 and 329 in the E326K mutant is driving GCase dimerization, we introduced negatively charged residues at position 329 (R329D or R329E) in combination with the E326K mutation and assessed the oligomeric state of the resulting proteins. We also designed another double mutant (E326K+R329A) as a negative control, hypothesizing that the uncharged alanine side chain would be unable to restore the stabilizing salt bridge with K326. For both the E326K+R329E and E326K+R329D mutants, SEC profiles revealed that introduction of a negatively charged residue at position 329 within the E326K backbone markedly shifts the elution profile from a higher-order peak (corresponding to the dimeric/oligomeric elution profile of the E326K variant alone) towards a smaller monomeric elution profile observed with WT GCase (Fig 3D). As expected, the E326K+R329A double mutant behaved as a constitutive dimer, with an elution profile similar to E326K. The monomeric and dimeric peaks were purified separately for the E326K+R329D and E326K+R329E double mutants, along with the dimeric species of the E326K+R329A double mutant. Biophysical characterization of the isolated monomeric and dimeric peaks revealed DLS and thermostability profiles comparable to those of WT and E326K proteins, respectively (Fig. S5B). All purified species of the double mutants were tested in both the 4-MU-β-Glc and liposome assays and showed comparable activity to imiglucerase, confirming that GCase oligomeric state does not impact its enzymatic activity under these experimental conditions (Fig. S5C, D).

We next sought to determine if restoration of the salt bridge between residues 326 and 329 was sufficient to restore the interaction between GCase and LIMP2 in cells using our co-immunoprecipitation assay. Introducing a negative charge at residue 329 restored LIMP2 binding, as both E326K+R329D and E326K+R329E mutants bound at levels comparable to WT GCase, whereas the E326K+R329A mutant showed similarly low binding as E326K alone (Fig. 3E, F). Collectively, these data provide structural rationale to explain how the E326K mutation leads to reduced lysosomal delivery of GCase and ultimately results in loss of lysosomal GCase activity by removing a key salt bridge that helps stabilize GCase in the monomeric state.

### GBA1-p.E326K cells accumulate GCase substrates and have enhanced lysosomal secondary lipid storage

To compare the extent of GCase substrate accumulation and secondary lipid storage observed between a PD-linked variant and GD-linked variant in *GBA1*, we analyzed total cellular and lysosomal lipid content from our Lyso-IP *GBA1* variant lines using LC-MS/MS. Significant accumulation of the GCase substrate GlcSph was present in E326K KI cells at both the whole cell and lysosomal level (Fig. 4A), and the extent of GlcSph accumulation within lysosomes correlated well with the degree of reduced lysosomal GCase activity and enzyme levels observed across all conditions assessed (Fig. 4B). Most GlcCer species were not significantly accumulated at the whole cell level except in the case of *GBA1* KO lines; however, relatively higher enrichment was observed for most GlcCers in isolated lysosomes (Fig. 4A), confirming that GCase substrates primarily accumulate within lysosomes. Surprisingly, broader lipidomic analysis of whole cell lysates and isolated lysosomes from E326K KI cells revealed aggravated perturbations to many lipid species compared to that observed in L444P KI and *GBA1* KO cells and lysosomes with near-total or total loss of GCase activity. When considering only specific lipids whose levels reached nominal significance (p > 0.1) in a genotype-level comparison of E326K cells and lysosomes to WT cells and lysosomes, we noticed that species from virtually every class of lipids in the glycosphingolipid (GSL) catabolic pathway that GCase functions in were represented (Fig. 4C, D & S6A, B). We also observed greater accumulation of specific BMP species in E326K lysosomes than in those isolated from cells with more severe GCase deficiency (Fig. 4E). Importantly, we showed that secondary lipid perturbations in E326K cells occur downstream of GCase LoF, as imiglucerase treatment fully corrected GlcSph (Fig. 4F) and partially normalized the levels of several secondary lipids (Fig. 4G, H). Our results confirm that the E326K variant triggers GCase substrate accumulation within lysosomes and reveal that this PD-linked variant leads to more pronounced secondary lipid storage compared to more severe GCase LoF.

**Figure 4.**
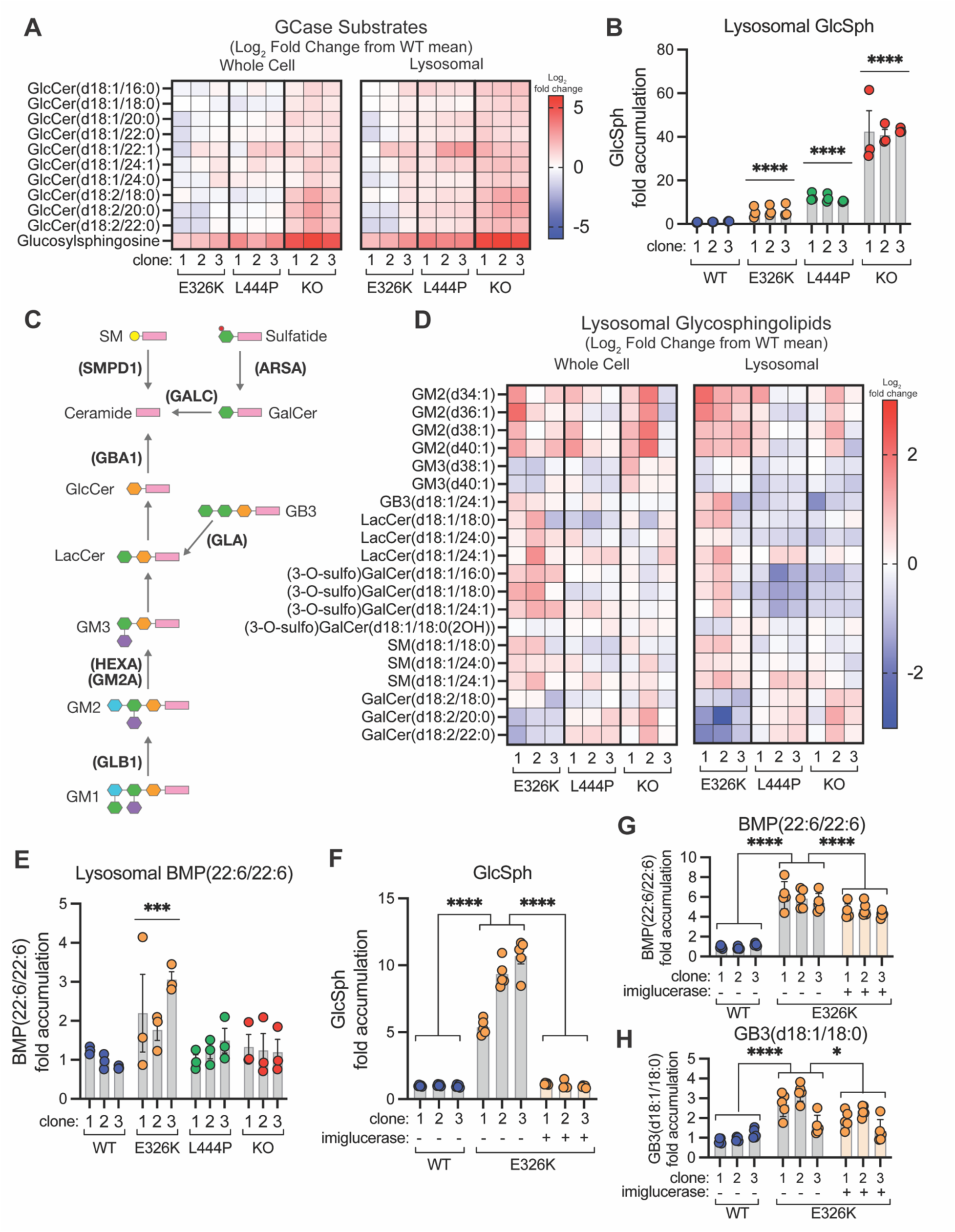
GBA1-p.E326K causes a mild accumulation of GCase substrates in the lysosome and greater perturbations to glycosphingolipid and BMP homeostasis. **A** Heatmaps depicting log_2_ fold changes (from WT mean) in the abundance of all detected GCase substrates from whole-cell and Lyso-IP fractions taken from WT, *GBA1*-p.E326K KI, *GBA1*-p.L444P KI, and *GBA1* KO HEK293T cells. **B** The fold change (over WT mean) of glucosylsphingosine (GlcSph) levels in Lyso-IP samples from HEK293T cells of the indicated genotype. **C** Schematic depicting the major lipids and key enzymes/protein co-factors in the lysosomal glycosphingolipid degradation pathway proximal to GCase. **D** Heatmaps showing the log_2_ fold change (from WT mean) of glycosphingolipids (from pathway shown in c) in both whole-cell and Lyso-IP fractions taken from HEK293T cells of indicated genotype. Only lipids with a fold change of E326K vs WT that reached nominal significance (p < 0.1) in either the whole-cell or Lyso-IP fractions are included. **E** Fold change (over WT mean) of BMP(22:6/22:6) levels in Lyso-IP samples from HEK293T cells of the indicated genotype. **F** Fold change (over WT mean) of glucosylsphingosine levels in whole cell samples from HEK293T cells of the indicated genotype, treated with imiglucerase as indicated. **G & H** Fold change (over WT mean) of BMP(22:6/22:6) (G) or GB3(d18:1/18:0) (H) levels in whole cell samples from HEK293T cells of the indicated genotype, treated with imiglucerase as indicated. All bar graphs show mean ± SEM with individual points representative of data from a single experimental replicate. Data in A-B, and D-E are from n = 4 independent replicates. Data in F-H are from n = 5 independent replicates. For A and D, statistical tests were performed on genotype-level comparisons using robust linear model as described in methods and significance threshold was set at FDR adjusted P value < 0.05. All other statistical tests were performed on genotype-level comparisons using one-way ANOVA with Dunnett’s multiple comparison test (B and E), or Tukey’s multiple comparisons test (F-H), where * = p < 0.0332, *** = p < 0.0002, **** = p < 0.0001.

*Lysosomal proteomics reveals functional consequences of GBA1-p.E326K-mediated dysfunction* To gain further insight into how the PD-linked E326K and GD-linked L444P variants compromise lysosomal function, we next performed unbiased proteomic analysis of lysosomes isolated from our suite of *GBA1* LoF cell lines. We found that lysosomes from E326K KI cells showed the greatest alterations in their protein composition compared to those observed in lysosomes from L444P KI or *GBA1* KO cells (Fig. 5A), with the number of differentially abundant proteins in lysosomes showing an inverse correlation with residual GCase activity across cell models. Intriguingly, this trend was opposite of what we observed in a proteomic analysis of whole cell lysates (Fig. S7A). Furthermore, we observed very little overlap in differentially abundant lysosomal proteins across genotypes (Fig. 5A), highlighting that each individual GCase variant, despite all inducing GCase LoF, has unique effects on lysosomal homeostasis. Pathway analysis revealed that proteins annotated as being mitochondrial^64^ in origin were among the most significantly and consistently depleted from E326K lysosomes compared to lysosomes isolated from L444P or GBA1 KO cells (Fig. 5B). Gene Ontology (GO) Cell Component analysis of significantly depleted proteins from lysosomes across genotypes revealed an overrepresentation of proteins associated with mitochondrial GO terms in E326K lysosomes, and very little overlap of proteins associated with these same terms was observed in either L444P or *GBA1* KO lysosomes (Fig. 5C).

**Figure 5.**
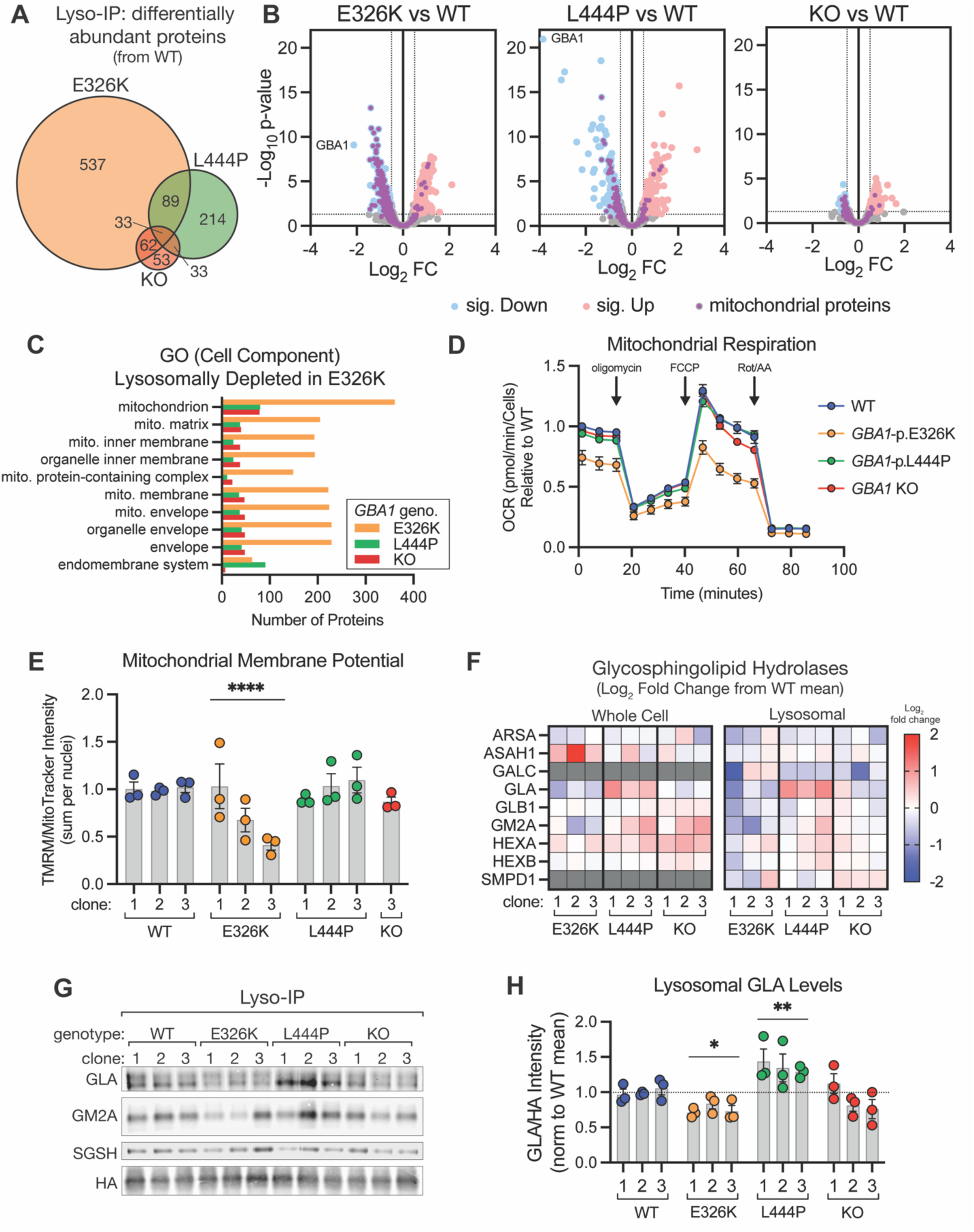
Proteomic analysis of purified lysosomes reveals GBA1-p.E326K specific cellular dysfunctions. **A** Venn diagram showing overlap in the identity of proteins with significantly altered abundance in lysosomes isolated from *GBA1*-p.E326K KI, *GBA1*-p.L444P KI HEK293T, and *GBA1* KO cells, relative to lysosomes isolated from WT cells. **B** Volcano plots showing differential abundance of proteins identified in Lyso-IP samples from indicated GBA1 variant or KO cells compared to WT cells. Proteins with significantly changed abundance (p < 0.05; absolute log_2_FC ≥ 0.5) are highlighted in red or blue for increased or decreased abundance, respectively. Proteins identified as mitochondrial in origin in the MitoCarta 3.0 database^64^ are highlighted in purple. **C** The number of proteins identified in each *GBA1* variant or KO genotype belonging to the top ten most significant GO: Cell Component terms associated with the subset of significantly depleted proteins in E326K (vs WT) lysosomes. **D** Oxygen consumption rate (OCR) of HEK293T cells of indicated genotype, measured using Seahorse XF Cell Mito Stress Test assay. n = 3 independent experiments. **E** HEK293T cells of indicated genotype were stained with TMRM and MitoTracker Green, and the ratio of the sum (per nuclei) TMRM fluorescence to MitoTracker Green fluorescence, normalized to the mean of all WT cells, is plotted. n = 3 independent experiments. **F** Heatmap of the relative abundance of all identified glycosphingolipid hydrolases (related to the pathway shown in 2c) is shown as log_2_ fold change (from WT mean) for both whole-cell and Lyso-IP samples from HEK293T cells of indicated genotype. Analytes that were not detected in a sample are shaded gray. **G** Representative immunoblot of Lyso-IP samples from HEK293T cells (of indicated genotype) expressing TMEM192-HA^x3^ tag. Immunoblots were probed with antibodies against GLA, GM2A, SGSH, and HA as a loading and lysosomal isolation control. **H** Quantification of lysosomal GLA levels (normalized per sample to HA band intensity), measured via immunoblotting of Lyso-IP fraction & normalized to mean of all WT samples. n = 3 independent experiments. Bar graphs in E and H show mean ± SEM with individual points representative of data from separate experimental replicates. All proteomics data (A-C, F) is from n = 3 independent experiments. Unless otherwise noted, all statistical tests were performed on genotype-level comparisons using robust linear model as described in methods and significance threshold was set at FDR adjusted P value < 0.05. Statistics in E and H were performed on genotype-level comparisons using one-way ANOVA with Dunnett’s multiple comparison test, where * = p < 0.0332, ** = p < 0.0021, **** = p < 0.0001.

Based on these results, we directly assayed mitochondrial function across our *GBA1* cellular models to understand how the PD- and GD-linked variants impacted mitochondrial homeostasis. We found significant deficits in mitochondrial respiration (Fig. 5D, S8A), as well as reduced mitochondrial membrane potential (Fig. 5E), specifically in E326K KI cells. In our whole cell proteomic analysis, we did not observe the same degree of mitochondrial protein depletion as we observed in purified E326K lysosomes (Fig. S7B), suggesting that these dysfunctional mitochondrial phenotypes are driven by changes at the lysosomal level. We next sought to explore how the E326K variant leads to mitochondrial protein depletion in lysosomes and defects in mitochondrial function. Impaired turnover of damaged mitochondria via mitophagy has been proposed to contribute to PD pathogenesis^65,66^, and we examined the impact of *GBA1*-p.E326K on the PTEN-induced putative kinase 1 (PINK1)/Parkin-dependent mitophagy pathway. We observed no difference in either the stabilization or activity of PINK1 or the turnover of select mitochondrial proteins in *GBA1* variant cells treated with CCCP overnight to induce mitochondrial damage (Fig. S8B-G), suggesting that the mechanism of GCase-dependent mitochondrial dysfunction appears unrelated to mitophagy.

Our proteomic analysis of lysosomal content from *GBA1* variant cells also yielded insights into the mechanisms underlying the enhanced secondary lipid storage phenotype in E326K lysosomes. In lysosomes from E326K KI cells, the levels of hydrolases (and related protein co-factors) involved in GSL catabolism (see Fig. 4C) were generally more perturbed than was observed in lysosomes from L444P or *GBA1* KO cells (Fig. 5F). While the overall effect size was mild, there was a remarkably consistent inverse correlation between GSL hydrolase levels in E326K lysosomes and levels of corresponding GSL species measured in E326K lysosomes via lipidomics (Fig. 4D, 5F). We confirmed key hydrolases were specifically depleted from E326K lysosomes compared to lysosomes isolated from WT cells by immunoblotting (Fig 5G, H & S7C-E). The broad dysregulation of GSL hydrolases in E326K lysosomes suggests that coordinated regulation of lysosomal composition controls GSL metabolism and that this balance is selectively disrupted by the trafficking defect of the E326K enzyme.

### GCase dysfunction and lipid storage phenotypes in murine brain and human CNS cell models of the E326K variant

Having demonstrated that *GBA1*-p.E326K has reduced endolysosomal GCase activity and triggers substrate accumulation in cellular models, we next wanted to explore whether this variant leads to similar changes in GCase pathway activity in the CNS. To assess this, we employed a mouse model in which the E326K variant has been knocked-in to the endogenous mouse *Gba1* locus. We analyzed GCase substrate levels in heterozygous and homozygous *Gba1* E326K KI mice and age-matched WT littermate controls by LC-MS/MS and observed a significant accumulation of GlcSph in brain lysates from these mice that was gene-variant dosage-dependent (Fig. 6A). We saw a significant reduction in GCase enzyme activity in brain lysates from homozygous *Gba1* E326K KI mice compared to WT mice, with a trend towards reduced activity also observed in brain lysates from heterozygous animals (Fig. 6B). This data correlated well with GCase enzyme levels measured in the same brain lysates via immunoblotting (Fig. 6C, D), suggesting a similar mechanism as we observed in our cellular models whereby loss of GCase protein underlies the GCase LoF phenotype. To assess the impact of the E326K variant on endolysosomal GCase activity in the mouse brain, we focused on microglia given their high expression of *Gba1* and strong dependence on lysosomal activity for their function^67–69^. We adapted a previously-described protocol for the dissociation and *ex vivo* analysis of mouse microglia^70^ and assessed GCase activity in this cell population using the LysoFQ-GBA probe. The results of this analysis showed a significant reduction in lysosomal GCase activity in microglia from homozygous E326K knock-in brains (Fig 6E), demonstrating that the E326K variant leads to a loss-of-function phenotype in the CNS.

**Figure 6.**
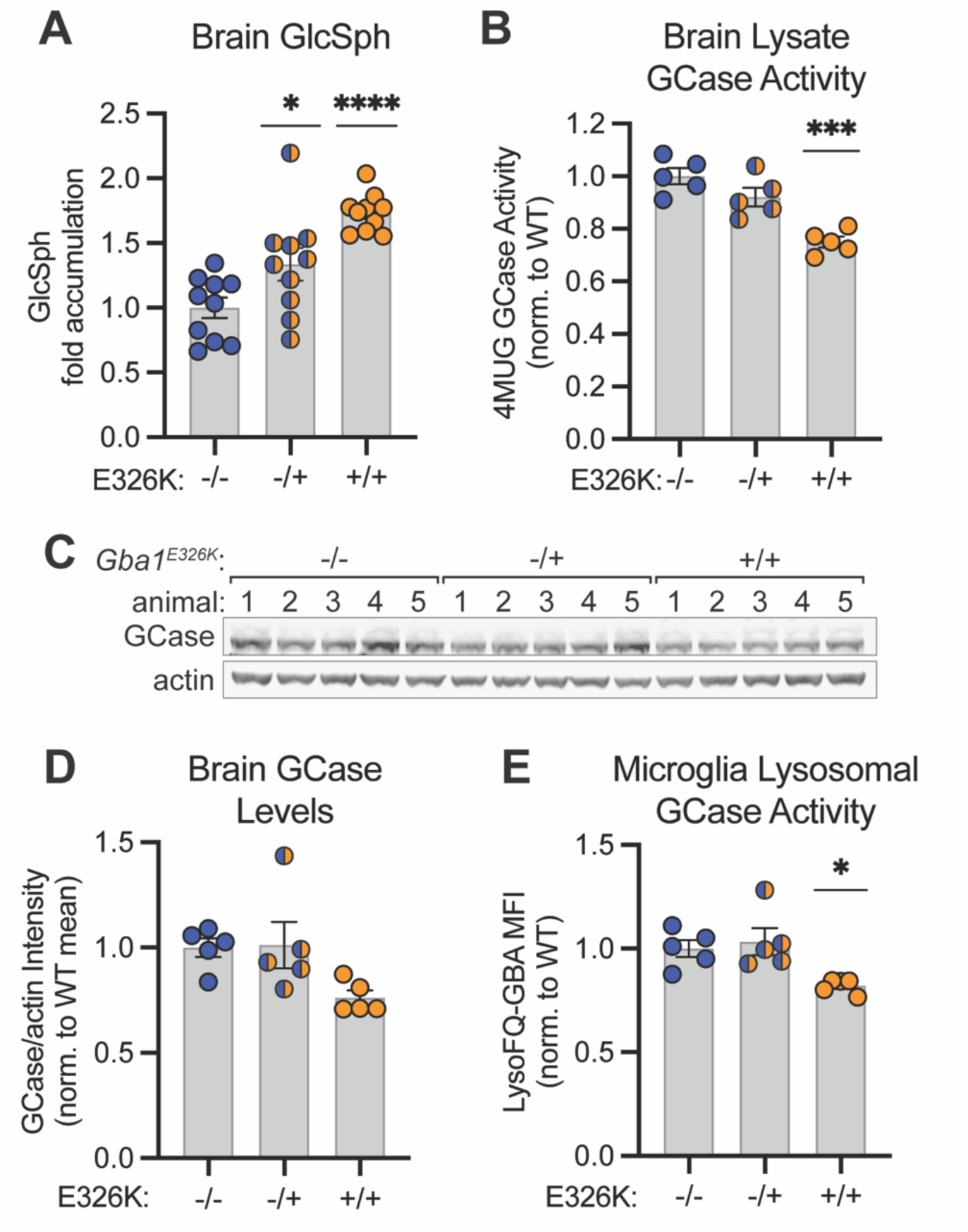
E326K knock-in mice recapitulate the primary GCase loss-of-function phenotypes observed in cellular models. **A** Glucosylsphingosine levels in bulk brain lysates from mice of indicated genotype are shown, normalized to the mean of all WT animals. n = 10 animals per genotype. **B** GCase enzymatic activity was measured in bulk brain lysates from mice of indicated genotype by assessing turnover of 4-MUG substrate. **C** Immunoblot showing GCase levels from bulk brain lysates from mice of indicated genotype. **D** Quantification of GCase band intensity (normalized per sample to actin band intensity), normalized to the mean of all WT samples. **E** Lysosomal GCase activity, quantified in microglial cell populations from dissociated mouse brain samples stained with LysoFQ-GBA. For B-E, n = 5 animals/genotype. All bar graphs depict mean ± SEM with individual points representative of data from an individual animal. Unless otherwise noted, all statistics were performed on genotype-level comparisons using one-way ANOVA with Dunnett’s multiple comparison test, where * = p < 0.0332, *** = p < 0.0002, **** = p < 0.0001.

Given our results from the mouse brain, along with emerging evidence implicating GCase dysfunction in immune system dysregulation in PD pathogenesis^71–74^, we next explored the impact of *GBA1*-p.E326K on GCase activity and glycosphingolipid homeostasis more broadly in human iPSC-derived microglia (iMicroglia). *GBA1*-p.E326K KI iMicroglia had reduced GCase activity and levels compared to WT cells (Fig. 7A-C), consistent with results from our HEK cell models. We next assessed the extent of GCase substrate accumulation in E326K KI and *GBA1* KO iMicroglia via LC-MS/MS. E326K KI iMicroglia significantly accumulated GlcSph, albeit to a lesser degree than that observed in *GBA1* KO iMicroglia (Fig. 7D, E). We also observed an accumulation of all GlcCer species measured in E326K KI iMicroglia and found that many GlcCer species were elevated to a greater extent than that measured in *GBA1* KO iMicroglia (Fig. 7D). To determine the impact of the E326K variant on GSL homeostasis in these cells, we performed targeted lipidomic analysis and measured the levels of GCase-relevant GSLs and BMPs in E326K KI and *GBA1* KO iMicroglia. E326K KI iMicroglia showed a striking secondary lipid accumulation phenotype in which the levels of all detected species of BMP and related metabolic precursors and many GSLs were elevated compared to WT and *GBA1* KO cells (Fig. 7F, G). In particular, ganglioside and globoside species were significantly elevated in E326K KI iMicroglia and showed a greater and, in some instances, an opposing effect to that observed in *GBA1* KO cells (Fig. 7G). Together, these results establish that the E326K variant leads to impaired GCase activity in a human CNS-relevant cell type and provide additional evidence that this variant triggers greater secondary lipid storage compared to more severe GCase loss-of-function.

**Figure 7.**
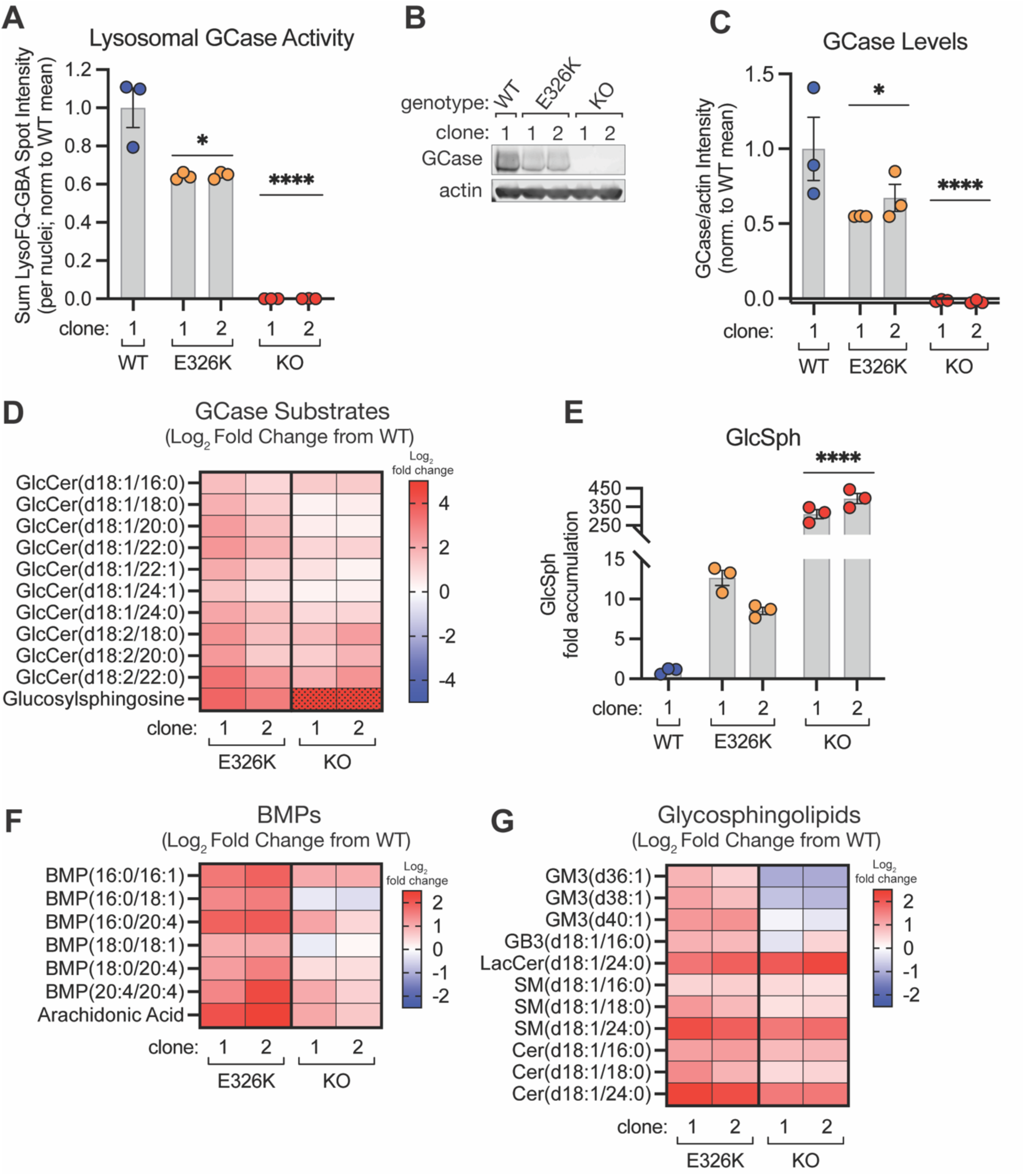
Human E326K-knock in iPSC-derived microglial cells have reduced GCase activity and accumulation of BMP and glycosphingolipids. **A** Lysosomal GCase activity, measured in WT, *GBA1*-p.E326K, and *GBA1* KO human iMicroglia stained with LysoFQ-GBA, is quantified as sum LysoFQ-GBA spot intensity per nuclei, normalized to the mean of all WT cells. n = 3 independent experiments. **B** Representative immunoblot showing GCase levels in human iMicroglia of indicated genotype. **C** Quantification of total cellular GCase enzyme levels (normalized to actin) in human iMicroglia of indicated genotype, measured via immunoblot & normalized to mean of all WT cells. n = 3 independent experiments. **D** Heatmap of whole-cell log_2_ fold change (from WT mean) abundance of all detected GCase substrates from human iMicroglia of indicated genotype. Shaded cells indicate values capped at log_2_ fold change = 5. **E** Fold change (over WT mean) of glucosylsphingosine levels in whole-cell samples from human iMicroglia of the indicated genotype. **F** Heatmap of log_2_ fold change (from WT mean) of BMPs & related species from human iMicroglia of indicated genotype. **G** Heatmap of log_2_ fold change (from WT mean) of glycosphingolipids from human iMicroglia of indicated genotype. For F and G, only species with a fold change of E326K vs WT that reached nominal significance (p < 0.1) are included. Lipidomics analysis (D-G) was done using a robust linear model as described in methods and significance threshold was set at FDR adjusted p value < 0.05, from n = 3 independent experiments. Statistics in A, C, and E, were performed on genotype-level comparisons using one-way ANOVA with Dunnett’s multiple comparison test, where * = p < 0.0332, **** = p < 0.0001.

### GCase LoF and lipid storage phenotypes identified in human GBA1-p.E326K variant carrier samples

Given the majority of *GBA1*-p.E326K PD patients carry a single variant allele, we next explored whether our findings on the consequences of the E326K variant on GCase activity and secondary lipid accumulation could be replicated in a heterozygous context. Our analysis of GCase activity measurements in whole blood samples from human subjects enrolled in the Parkinson’s Progression Markers Initiative (PPMI) confirmed recent data reporting that GCase activity is significantly reduced in E326K carriers (irrespective of PD diagnosis) compared to individuals with no known *GBA1* mutation (Fig. 8A)^75^. We recently completed a broad lipidomic assessment of the levels of relevant lipids in plasma samples from a similar PPMI-based cohort of individuals (Table S3, S4), and we observed a significant increase in GlcSph levels in plasma from heterozygous E326K carriers compared to individuals that do not carry mutations in *GBA1* (Fig. 8B). Consistent with our results from cellular models, the extent of impaired GCase activity and GlcSph accumulation was lower in E326K carriers compared to carriers of a *GBA1* mutation classified as “pathogenic” (predominantly the N370S variant in this cohort) (Fig 8A, B). We performed a high-level analysis to identify lipids that were specifically dysregulated in E326K carriers and were intrigued by the observation that the globoside GB3(d18:1/24:0), a GSL in the same catabolic pathway that GCase functions in, was significantly accumulated specifically in plasma from E326K carriers compared to non-*GBA1* variant subjects (Fig. 8C). Given the inter-subject heterogeneity in human patient samples, especially for secondary phenotypes downstream of a primary genetic perturbation, this data is remarkably consistent with the enhanced secondary lipid storage we observed in cellular models. Moreover, these results suggest that key mechanisms of both GCase and lysosomal dysfunction caused by the E326K variant are relevant in the context of heterozygous expression in human subjects.

**Figure 8.**
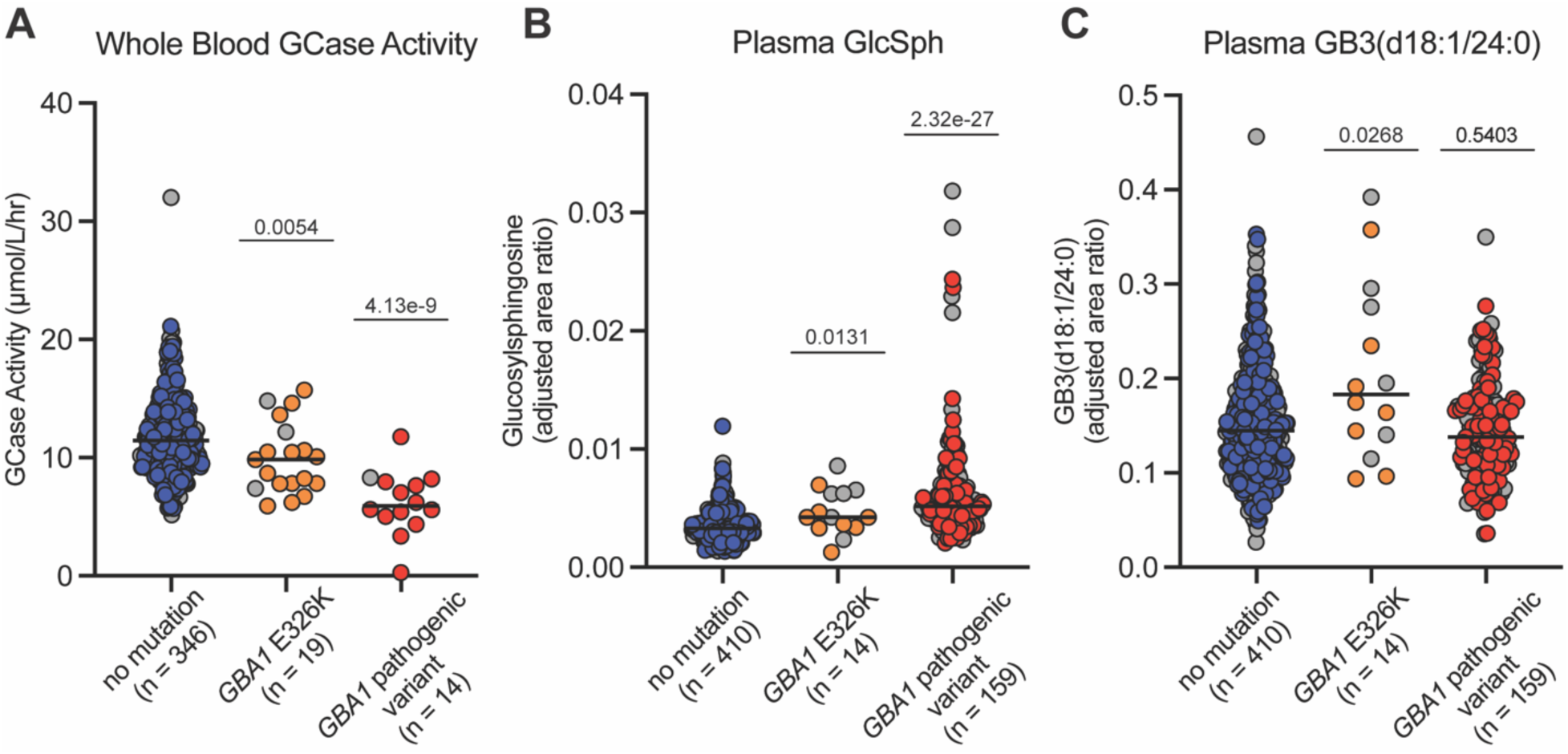
Evidence of GCase loss-of-function and enhanced secondary lipid storage in biofluid samples from human *GBA1*-p.E326K carriers in the PPMI cohort. **A** Whole blood-derived GCase activity (µmol/L/hr) is plotted in *GBA1*-p.E326K carriers (n = 19), *GBA1* pathogenic variant carriers (n = 14), and subjects who do not carry either (n = 346). **B** and **C** The levels of selected plasma metabolites (adjusted area ratio), glucosylsphingosine (**B**) and GB3 (d18:1/24:0) (**C**), plotted from *GBA1*-p.E326K carriers (n = 14), *GBA1* pathogenic variant carriers (n = 159), and subjects who do not carry either (n = 410). P-values are derived from linear models comparing *GBA1*-p.E326K carriers or *GBA1* pathogenic variant carriers to non-carriers, adjusted for age, sex, PD case status, and 3 principal components derived from genome-wide WGS data (see Methods). Gray circles denote controls and colored circles (blue, orange, or red, for non-carriers, p.E326K carriers, and pathogenic variant carriers, respectively) denote PD cases.

## Discussion

While many variants in *GBA1* are linked to both Gaucher disease and PD, the specific association of the common E326K variant with risk for developing PD, but not GD, has remained enigmatic. Our study demonstrates that the E326K variant causes GCase LoF that is milder than that observed with the GD- and PD-linked variant L444P and arises from reduced delivery of GCase to lysosomes. Through biophysical and biochemical studies of recombinant E326K, we discovered that this mutation induces constitutive dimerization of the enzyme without impacting its intrinsic enzymatic activity. Structural analysis of the E326K variant, both alone and in complex with LIMP2, reveals that this variant results in the loss of a salt bridge that both enhances the dimerization of the enzyme and impairs its ability to interact with LIMP2 in cells. These structural alterations ultimately result in a reduction in lysosomal GCase enzyme levels that is accompanied by a reduction in lysosomal GCase activity and accumulation of the GCase substrate GlcSph. Surprisingly, we observe that E326K variant cells exhibit additional alterations to lysosomal and mitochondrial homeostasis that do not manifest in cell models with more severe GCase LoF. Analysis of human biofluid samples from *GBA1*-p.E326K carriers confirmed this primary LoF phenotype, as well as providing evidence for broader secondary lipid dysregulation in carriers of the E326K variant. As both lysosomal and mitochondrial dysfunction have been linked to PD pathogenesis^66,76,77^, these results raise the possibility that it is the combined lysosomal and mitochondrial dysfunction induced by the E326K variant that may help to explain the specificity of this variant’s association with PD and not GD.

Our results provide definitive and conclusive evidence that the E326K variant causes a loss of lysosomal GCase activity in cells, at least in part driven by impaired trafficking of the mutant enzyme to the lysosome. Lipidomic and proteomic profiling of lysosomal content from E326K variant cells reveals more extensive depletion of GCase enzyme and greater accumulation of GCase substrates than observed at the whole cell level, and our analysis of glycosylation patterns present on endogenous E326K variant enzyme are consistent with impaired trafficking of the enzyme to the lysosome. Prior work in GD patient-derived fibroblasts has demonstrated that for the two most prevalent GD-associated variants, ER retention and ERAD-mediated degradation results in reduced lysosomal GCase enzyme levels^4,5^. Of note, when compared to isogenic cell models harboring the L444P variant, the GCase dysfunction observed in the E326K variant cells is mild, perhaps explaining the lack of association between this variant and GD. Interestingly, GWAS studies have also identified SNPs in the *SCARB2* locus (which encodes LIMP2) with increased risk and earlier age of onset for PD^29,78,79^. LIMP2 has also been identified as a genetic modifier of PD-like pathologies^80^, and LIMP2-dependent trafficking of GCase has been shown to be defective in idiopathic PD patient-derived cells^81^, suggesting that reduced lysosomal delivery of GCase may be a common mechanism for disease risk and/or progression. The question of whether other PD-specific *GBA1* variants impact GCase activity through similar mechanisms warrants future study. While other studies have not documented signs of protein misfolding or ER retention with the E326K variant^16^, we demonstrate here that when overexpressed in cells, the E326K variant enzyme interacts less efficiently with its trafficking receptor LIMP2 than the WT enzyme does, providing mechanistic insights into the nature of the observed GCase LoF phenotype.

Structural and biophysical studies, including both crystallography and cryoEM, revealed that the E326K variant forms a constitutive dimer that remains stable across both neutral and acidic pH conditions, in contrast to WT GCase which is largely monomeric. While higher order species of GCase have been observed in material isolated from human tissue^82,83^ and dimerization has been previously reported and suggested to be functionally important, evidence of GCase homodimers could only be captured at high concentration in solution^59^ or in presence of small molecules^58^.

Prior studies have reported that maintaining a dynamic dimer-monomer distribution is important for promoting GCase activity in the lysosome, and the constitutive dimerization observed here with the E326K variant would be predicted to be associated with reduced lysosomal GCase activity as robust dimerization may inhibit complex dissolution and activation by SapC^57,59,84^.

Our cell biology studies do not preclude this possibility, but suggest that the primary mechanism by which the E326K variant reduces lysosomal GCase activity occurs upstream of its interaction with SapC. Overall, the organization of the two E326K dimers observed in this work is reminiscent of dimers observed in previously solved crystal structures, in which the active sites of both protomers are facing each other^57,58^. The catalytic site does not appear to be impacted by the E326K variant, consistent with our biochemical data showing no differences in the catalytic activity of E326K compared to recombinant WT enzyme. While the E326K mutation itself is not part of the dimer interface, a key finding from our structural studies is the discovery that a salt bridge between the side chains of residues E326 and R329 is disrupted by the E326K mutation. This subtle structural change appears to regulate the multimeric state of the enzyme as well as its interaction with LIMP2, as substitution of a negatively charged acidic residue (D or E) at position 329 shifts the equilibrium of recombinant E326K variant enzyme back towards monomeric species and also largely rescues the interaction with LIMP2 in cells. Notably, this mechanism is not unique to the E326K variant as several genetic mutations in other proteins linked to a wide range of diseases have been shown to alter multimeric assembly resulting in protein destabilization^85–88^, including in other neurodegenerative diseases^89,90^. Overall, our findings represent a striking example of a GCase variant adopting a stable multimeric assembly. Additional studies are warranted to determine whether other PD-specific *GBA1* variants also impact GCase activity by modulating its oligomeric state.

Our *in vitro* characterization and cellular studies suggest that impaired interactions with LIMP2 likely underpin the defective lysosomal trafficking observed with the E326K variant. Our binding and structural studies demonstrate that, while the E326K variant remains competent to interact with LIMP2, forming a stable 2:2 complex, the affinity with which the E326K dimer binds LIMP2 at neutral pH is lower compared to the affinity of WT monomer for LIMP2 under identical conditions. Intriguingly, recent molecular dynamic modeling of the E326K variant suggests that this mutation may alter the stability of the GCase dimer^91^. While this same report predicts increased affinity between the E326K variant and LIMP2, our data are not wholly inconsistent with the premise that this mutation alters the multimeric state of the enzyme in a manner that affects its trafficking to the lysosome. Our co-immunoprecipitation studies using cell lysates showed that the interaction between LIMP2 and the E326K variant is significantly impaired, suggesting a greater magnitude of defect in binding than that observed in our *in vitro* analyses. Several possibilities exist to explain the apparent discrepancy we observe between the strength of the interaction between the E326K variant and LIMP2 *in vitro* and in cells. First, our studies using recombinant LIMP2 utilize only the lumenal domain of the protein, and structural constraints imposed by the N- and C-terminal transmembrane helices of LIMP2 embedded in a membrane may not accommodate efficient binding to an E326K dimer in cells. It is also possible that the dimeric organization observed in the recombinant system does not reflect the native state in E326K, with dimerization instead acting as a compensatory mechanism to stabilize the mutant enzyme under overexpression conditions. Indeed, while other studies have described similar binding impairment of the E326K variant to LIMP2, preliminary characterization of the recombinant complex in those studies did not suggest heterotetrameric assembly^49^. Additional interactions with factors absent from our *in vitro* system, such as other proteins or biological macromolecules, may also contribute to the differences in binding between GCase E326K and LIMP2 in cells. Whether or not the multimeric organization of the E326K variant is directly related to the mechanism by which E326K variant enzyme is less efficiently trafficked to the lysosome, or if this perceived reduction in binding to LIMP2 in cells results from the destabilization of the variant enzyme, is an important area for future investigation.

Across cell models employed in this study, glycosphingolipid (GSL) metabolism appears disproportionately perturbed in E326K variant cells with respect to the severity of the GCase LoF phenotype in these cells. Lipidomic profiling of E326K variant cells and purified lysosomes revealed alterations to virtually every class of GSLs metabolically linked to GCase and many species of BMPs to a degree not observed in L444P KI or *GBA1* KO cells that have more severely compromised GCase function. Proteomic analysis of lysosomal content also revealed inverse correlation in E326K variant lysosomes between GSL abundance and many of the hydrolases and cofactors involved in their catabolism in the lysosome, hinting at possible mechanism by which coordinated trafficking of lumenal proteins regulates lysosomal metabolism by affecting lysosomal composition. It is intriguing to speculate that this disconnect between severity of GCase dysfunction and secondary lipid dysregulation in E326K variant cells might be related to compensatory mechanisms that are triggered in the case of severe lysosomal dysfunction, which may not be fully engaged in the case of a mild LoF mutation like E326K. As lysosomes function not only as key mediators of cellular catabolism, but also as signaling platforms and sites of ion and nutrient storage^92,93^, cells invest significant resources in ensuring lysosomes function properly, and a myriad of pathways have been characterized that regulate lysosomal biogenesis^94^, positioning and function^95^, and maintenance of membrane integrity.^96^ Given the severity of GD in comparison to PD, it is possible that chronic, low-level GCase dysfunction observed with the E326K variant manifests additional or exacerbated lysosomal pathologies over time, while severe GCase dysfunction may trigger a protective, but ultimately futile, compensatory response. ^64,98^ Given that LIMP2 has been reported to regulate cellular and lysosomal lipid homeostasis independently of GCase ^61,97^, the secondary lipid storage in E326K cells could also arise from impaired LIMP2 function downstream of altered interactions with the variant enzyme.

Lysosomal dysfunction has broad consequences for cellular homeostasis, and emerging evidence has highlighted the importance of functional connections between lysosomes and mitochondria, especially in the context of neurodegenerative disease such as PD.^98–105^ GCase itself has been functionally linked to mitochondria via multiple mechanisms. Impaired GCase function has broadly been reported to impact autophagy and the clearance of defective mitochondria^106,107^, mitochondria-lysosome contacts have been reported to be dysregulated in cells harboring GCase loss-of-function variants^99^, and GCase has also been reported as a regulator of mitochondrial respiration and energy metabolism.^108^ Our results also suggest a functional connection between GCase and mitochondria – perhaps the most surprising feature of our proteomic profiling of lysosomes from *GBA1* variant cells is the observation that proteins of mitochondrial origin appear depleted more significantly from E326K variant lysosomes. While it is not immediately clear how this phenotype is mechanistically related to mitochondrial function, we observe functional deficits in mitochondria specifically in E326K cells that suggests a defect in the crosstalk between mitochondria and lysosomes. Furthermore, we observe no gross defects in the turnover of select mitochondrial proteins, nor the induction of PINK1 activity, in E326K cells where we pharmacologically depolarized mitochondria, suggesting that E326K-associated mitochondrial dysfunction is not intrinsic to mitochondria, but instead is downstream of lysosomal dysfunction. It is tempting to speculate that mitochondrial dysfunction may be related to the enhanced secondary lipid accumulation observed in E326K lysosomes, especially considering the observation that mitochondrial dysfunction in other LSDs affecting GSL catabolism.^103^ Further studies are warranted to explore both the mechanisms underlying the association of this variant with mitochondrial function and the potential impact of *GBA1*-associated mitochondrial dysfunction on PD pathogenesis.

Our studies demonstrate that the GCase LoF and secondary lipid dysregulation caused by the E326K variant occurs in relevant CNS cells and human patient samples. In both human iPSC-derived microglia and *ex vivo* mouse microglia, we show that the E326K variant results in a loss of GCase enzyme and induces accumulation of GCase substrates. Macrophages are the primary peripheral cell type affected by GCase loss-of-function in GD, and thus we were particularly interested to see strong phenotypes stemming from GCase dysfunction in a myeloid-lineage CNS cell type like microglia.^72^ Additionally, a recent report characterizing an E326K knock-in mouse highlighted increased neuroinflammation and microgliosis in both the substantia nigra and dentate gyrus at baseline in knock-in mice.^17^ Importantly, these findings of GCase dysfunction & substrate accumulation were also replicated in an analysis of human biofluid samples from heterozygous carriers of the E326K variant. Our analysis of human plasma samples suggests that enhanced secondary lipid accumulation, similar to what we observe in homozygous E326K cell models, also occurs in *GBA1*-p.E326K heterozygous carriers. These observations provide evidence that a single copy of the E326K variant is sufficient to elicit phenotypes in humans that are indicative of GCase dysfunction, and increased risk for PD. Currently, numerous strategies are being explored as disease-modifying therapies for the treatment of *GBA1*-PD. Collectively, our analysis of the mechanisms of dysfunction stemming from a common PD-associated *GBA1* variant highlights the potential for therapeutic approaches aimed at increasing GCase enzyme levels and identifies potential biomarkers for monitoring lysosomal pathway correction with *GBA1*-targeted therapies.

## Methods

### Antibodies

Primary antibodies used include:

GCase (rb mAb, abcam, ab125065; 1/1000)

β-actin (ms mAb, Sigma-Aldrich, A2228; 1/2000)

HA-Tag (rb mAb, Cell Signaling Technology, #3724; 1/2000)

NPC2 (rb pAb, Proteintech, 19888-1-AP; 1/1000)

LIMP2 (rb mAb, Thermo, 703037; 1/1000)

Strep-tag (ms mAb, Qiagen, 34850; 1/250)

GLA (ms mAb, Proteintech, 66121-1-Ig; 1/1000)

GM2A (rb mAb, abcam, ab313587; 1/1000)

SGSH (rb mAb, abcam, ab200346; 1/1000)

GCase (rb pAb, Sigma-Aldrich, G4171; 1/1000)

phospho-Ubiquitin (Ser65) (rb mAb, Cell Signaling Technology, #62802; 1/1000)

PINK1 (rb mAb, Cell Signaling Technology, #6946; 1/1000)

TIM23 (ms mAb, BD Biosciences, 611222; 1/1000)

PDHA1 (ms mAb, abcam, ab110330; 1/1000)

Mitofilin (ms mAb, Proteintech, 68226-1-Ig; 1/1000)

### HEK293 Cell Culture & Generation of GBA1 Variant or Overexpressing Cell Lines

Unless otherwise noted, all HEK293 cultures were maintained in a complete culture medium comprised of DMEM (Thermo 11965092) supplemented with 10% fetal bovine serum (VWR 97068-085)), in a 37°C humidified atmosphere with 5% CO_2_.

Gene-edited HEK293 lines were generated as follows: for GBA1 KO a guide sequence 5’-GCCCGTGTGATTAGCCTGGATG-3’ targeting GBA1 was assembled with a sgRNA scaffold and inserted into the sgRNA cloning site of pRSGCCG-U6-sg-CMV-Cas9-2A-TagGFP2 (Cellecta SVCRU6CCG-L) per the manufacturer’s instructions. HEK293 with the E326K or L444P variants knocked into the endogenous GBA1 locus were generated as follows: guide sequences 5’-CAGGCGGTGTGTCTCCCCTA-3’ (for E326K) or 5’-TGCCAGTCAGAAGAACGACC-3’ (for L444P) were inserted into the sgRNA cloning site of pCas-Guide-EF1a-GFP (Origene GE100018) per the manufacturer’s instructions. Corresponding ssDNA templates (synthesized at IDT) were used as homology donor templates to knock in the mutation of interest: for E326K the sequence 5’-CTAAATATGTTCATGGCATTGCTGTACATTGGTACCTGGACTTTCTGGCTCCAGCCAAAGCCACGCTAGGGGAGACACACCGCCTGTTCCCCAACACCATGCTCTTTGCCTCAGAGGCCTGTGTGGGCTCCAAG-3’ was used, and for L444P the sequence 5’-GCAAGTTCATTCCTGAGGGCTCCCAGAGAGTGGGGCTGGTTGCCAGTCAGAAGAACGACCCAGACGCCGTGGCACTGATGCATCCCGATGGCTCTGCTGTTGTGGTCGTGCTAAACCGGTGAGGGCAA-3’ was used. WT HEK293 cells (ATCC CRL-1573) were transfected with the resulting plasmid (and matched ssDNA donor template for knock-in mutations), and individual clones were isolated by single cell sorting of GFP+ cells by FACS. Homozygous GBA1 KO or GBA1 variant KI was confirmed by sanger sequencing genomic DNA extracted from individual clonal populations.

HEK293 cells expression the TMEM192-HA^x3^ Lyso-IP construct were generated via lentiviral transduction as described in Wang et al 2023. Cell cultures were maintained in complete medium supplemented with 200ug/mL Hygromycin B (Thermo 10687010) to maintain transgene expression, although Hygromycin B was not included in cultures prepared for experimental analysis.

### Cell Treatments

For imiglucerase rescue experiments, Hek293 cells were cultured on poly-L-lysine treated tissue culture dishes and maintained in complete culture medium. For measurement of lipids by LC-MS/MS WT and E326K cells were plated at a density of 60K per well in a six well plate on day zero. On day one cells were treated by replacement of the entire medium with fresh medium with or without addition of 2µM imiglucerase (Sanofi). Medium was replaced on days 2 and 3 (24 and 48h post addition). Cells were harvested on day 4 (72h post addition) for lipidomic analysis as described below.

For mitophagy induction experiments, cells were plated in dishes pre-coated with Poly-L-Lysine, and then treated with 10µM CCCP (Sigma, C2759) for 16 h, before harvesting for immunoblotting as detailed below.

### Preparation of cell extracts for LC-MS/MS

Cells were harvested by aspirating medium then briefly washed with 1mL of ice-cold 0.9% saline solution. Cells were extracted by scraping directly into 400uL ice-cold extraction buffer containing internal standards. Extraction buffer 9:1 v/v LSMS grade MeOH/Ultrapure Water containing 2uL internal standards per sample (ie: 2uL I.S./400uL). Extracts were transferred into 1.5mL Eppendorf tubes and shaken at 2000 r.p.m for 20 minutes followed by centrifugation at 21K x G at 4C for 5 minutes. Cleared supernatant was transferred to LCMS glass vials then dried under N2 gas then stored at –80C until analysis.

### LysoFQ GCase activity assay

LysoFQ-GBA staining of HEK cells was performed following previously published protocols (Deen, et al. 2022), and staining parameters were optimized in HEK cells prior to quantitative analysis using WT and GBA KO cells to ensure that the probe concentration and stain length used yielded results in the linear range of the probe turnover. Briefly, cells were seeded in the inner 60 wells of PDL-coated 96-well imaging plates (Corning, 354640) the day prior to staining in normal culture media. For analysis of iMicroglia cultures, plates were additionally coated with N-terminus truncated Vitronectin (rhVTN-N, Fisher A14700, 50ug/mL final) for 1h. As a positive control for the probe function, one well per cell line was pre-treated with 100µM conduritol-b-epoxide (CBE, abcam, ab144118) for 1h prior to staining. Cells were stained in complete culture medium, supplemented with 10µM LysoFQ-GBA probe and 2µM Hoechst 33342 (Thermo 62249), for 1h at 37°C. Cells were washed 3x with 1xEBSS (Thermo 14155063) and then incubated in a solution of 1xEBSS supplemented with 5% FBS, 5.55mM D-glucose, and 10µM isofagomine D-tartrate (Cayman Chemicals, 16137) to quench probe turnover for 15 minutes prior to imaging. Cell plates were imaged on an automated confocal high-content imager (Revvity, Opera Phenix High-Content Screening System) using a 40x water immersion objective lens, 405nm and 488nm excitation lasers, and preset emission filters. Channel acquisition was separated to avoid fluorescence spillover between channels. A custom analysis in the Harmony 4.9 software (Revvity) was used to quantify LysoFQ-GBA signal from the resulting images. Cell nuclei were segmented and counted from Hoechst stain using the “Find Nuclei” building block, and LysoFQ-GBA signal was segmented using the “Find Spots” building block. Lysosomal GCase activity was quantified as the sum corrected LysoFQ-GBA spot intensity was normalized to the number of nuclei in each image, and the average Lysosomal GCase activity per well was calculated from 12 fields per well. A total of four wells per individual cell line were imaged per plate, and the average Lysosomal GCase activity across all four wells is reported for a single experimental replicate.

### Recombinant protein purification

A DNA fragment coding for residues A40 to Q536 of human GBA1 was synthesized (Genscript), codon optimized for mammalian expression and cloned into a derived pRK5 vector containing a mouse kappa light chain signal sequence at its N-terminal and an uncleavable C-terminal 6x histidine tag. Protein was expressed in ExpiCHO cells for 7 days. Cells were harvested by centrifugation at 3000rpm for 10min, supernatant was filtered through a 0.22 µM membrane and loaded onto a Ni-NTA Xcel resin pre-equilibrated with 25mM HEPES pH 7.5, 150mM NaCl (Buffer A); resin was washed with Buffer A supplemented with 15mM imidazole and protein was eluted with Buffer A with 300mM imidazole. Elution fraction was concentrated down using an Amicon Ultra Centrifugal Filter (30 kDa MWCO) and delipidated by addition of 20% (v/v) butanol; reaction was nutated for 1h at room temperature and spun down at 10k rpm for 10 minutes. Aqueous fraction was collected and loaded onto a Superdex S200 16/60 pre-equilibrated with 20mM sodium acetate pH 5.5 and 150mM NaCl. Fractions containing protein were analyzed by SDS-PAGE and pooled. E326K & R329 mutations were sequentially introduced using a Quikchange site-directed mutagenesis into the wild-type backbone and expression and purification of resulting variants were carried out as described above.

A DNA fragment coding for the extracellular domain of human LIMP2 (residues S36 to N430) was synthesized (Genscript), codon optimized for mammalian expression and cloned into a derived pRK5 vector containing a mouse kappa light chain signal sequence at its N-terminal followed by a 6x histidine tag, Avitag and TEV protease cleavage site. Protein was expressed in HEK293 cells for 5 days. Cells were harvested by centrifugation at 3000rpm for 10min, supernatant was filtered through a 0.22 µM membrane and loaded onto a Ni-NTA Xcel resin pre-equilibrated with 25mM HEPES pH 7.5, 150mM NaCl (Buffer A); resin was washed with Buffer A supplemented with 15mM imidazole and protein was eluted with Buffer A with 300mM imidazole. Elution fraction was concentrated down to 5mL using an Amicon Ultra Centrifugal Filter (30 kDa MWCO) and loaded onto a Superdex S200 16/60 pre-equilibrated with 20mM sodium acetate pH 5.5 and 150mM NaCl. Fractions containing protein were analyzed by SDS-PAGE and pooled.

### In-vitro biotinylation

Purified proteins were biotinylated using a BirA biotin-protein ligase standard reaction kit from Avidity LLC following vendor’s recommendation. Once reactions reached completion, biotinylated proteins were polished over S200 as described above.

### Surface Plasmon Resonance (SPR) Binding Measurements

The surface plasmon resonance experiments were performed using a Biacore8K SPR System equipped with a Series S Sensor Chip SA. The Sensor Chip SA was equilibrated overnight (12hrs) at the pH condition of pH 5.5 (20mM Na acetate pH 5.5 + 150mM NaCl + 0.1% BSA + 0.005% Tween20) or pH 7.5 (25mM HEPES pH 7.5 + 150mM NaCl + 0.1% BSA + 0.005% Tween20). Biotinylated hLIMP2 was diluted in the corresponding pH condition and immobilized as the ligand at a density of 80-95RU. GCase analytes were diluted in corresponding buffers and injected in both pH conditions at concentrations of 0.8, 4, 20, 100 and 500 nM at a flow rate of 30 μl/min and at a temperature of 25°C in a single-cycle kinetic format. An association time of 300s was utilized, with a final dissociation of 600s. Each condition was repeated in duplicate channels, with an entire experimental replicate further confirming results. Data were collected at a rate of 10 Hz. The data were fit to a 1:1 Langmuir interaction model within the Biacore Insight Evaluation Software, with the exception of E326K as the analyte which was fit with a bivalent model. For modeling the bivalent fit, analyte concentrations reflect the E326K dimer concentration.

### Differential Scanning Fluorimetry

Melting experiments were conducted on a Prometheus NT48 (NanoTemper technologies) by measuring the tryptophan fluorescence 330/350 nm ratio of protein samples concentrated at 1 mg/mL in a standard capillary.

### Dynamic Light Scattering

DLS experiments were conducted on a Prometheus Panta (NanoTemper technologies) using samples concentrated at 1 mg/mL in a standard capillary.

### SEC-MALS analysis

GCase E326K/LIMP2 complex was run on a Acquity Premier SEC column (250Å, 1.7μM, 4.6 x 150mm, Waters) column equilibrated with 25mM HEPES pH 7.5 and 150mM NaCl in line with a Dawn HELEOS II (Wyatt Technologies) light scattering detector connected to a Wyatt OptiLab rEX refractive index detector. Wyatts Technologies software (ASTRA) was used to determine the molecular weight of the complex based on refractive index.

### Crystallization of GCase E326Kvariant

Protein formulated in 20mM sodium acetate pH 5.5 and 150mM NaCl was concentrated to 20 mg/mL and set-up intro crystallization trials. Crystals were grown at 19 °C by the hanging-drop method by mixing the resulting reaction with an equal volume of reservoir solution containing 0.09 M Sodium nitrate, 0.09 Sodium phosphate dibasic, 0.09M Ammonium sulfate, 0.1 M HEPES / MOPS pH 7.5 and 12.5% MPD, 12.5 PEG 1000 (w/v), 12.5% (w/v) PEG 3350 (Morpheus HT screen, condition C8 from Molecular Dimensions MD1-47). Harvested crystals were flash frozen in liquid nitrogen.

### X-ray data collection and structure determination

X-ray diffraction data were collected at EMBL Hamburg beamline P13^109^. Data was processed and scaled to a resolution of 3.1 Å using DIALS and Xia2^110,111^. Phases were obtained through molecular replacement using Phaser with chain A from PDB 2NT1 as the search model. Models were iteratively rebuilt and refined using COOT and REFMAC5^112,113^. Figures were generated using PyMOL^114^ and ChimeraX^115^.

### CryoEM sample preparation and data acquisition

Recombinant GCase E326K was incubated with a 1:2 molar excess of LIMP2 and incubated on ice for 30 min. The sample was injected over a Superose 6 3.2/300 GL column (GE Healthcare) equilibrated in 25 mM HEPES pH 7.5, 150 mM NaCl and peak fractions were collected. Samples from the peak fractions (∼3.5µl) were applied to glow discharged Quantifoil R1.2/1.3 Au300 holey carbon grids and subsequently flash frozen in a liquid ethane/propane (40/60) mixture using a Vitrobot (Thermo Fisher) set at 4°C and 100% humidity with a blot time range from 3 to 4.5 s.

Images were collected using a 200 keV Glacios (Thermo Fisher Scientific) with a Falcon 4 direct electron detector at a nominal magnification of 120,000x and calibrated pixel size of 1.2 Å/px. Movies were automatically collected using SerialEM^116^ using a multishot array. Data were collected at an exposure dose rate of ∼15 electrons/pixel/second as recorded from counting mode. Images were recorded for ∼2.7 seconds in 60 subframes to give a total exposure dose of ∼50 electrons per Å.

### CryoEM image processing

Image processing was performed in CryoSPARC v4.4.1^117^, as summarized in Fig. S3. After Patch Motion Correction and Patch CTF Estimation, exposures with CTF fit resolutions worse than 3.5 Å were discarded. Particles were selected using the blob picker (min diameter = 50 Å, max diameter = 100 Å) and extracted with a box size of 324 px and binned to 128 px. The particles were subjected to 2D classification, and the best 2D classes were used to generate an initial model via ab initio reconstruction. This model was used to generate templates for template picking. Multiple rounds of 2D classification were used to remove obvious junk particles. The remaining particles were subjected to iterative rounds of multi-class ab initio reconstruction and heterogeneous refinement. The particles in the best 3D class were re-extracted with a box size of 324 px and used for non-uniform refinement with C2 symmetry to generate a reconstruction with a GS-FSC resolution of 3.3 Å. Symmetry expansion was applied to the final particle stack prior to local refinement, leading to a final reconstruction with an overall resolution of 3.2 Å. Crystal structures of GCase (PDB: 2NT1, chain A) and LIMP2 (PDB: 4TW2, chain A) were used as initial models and docked into the map using ChimeraX. The model was then rebuilt and refined using COOT, ISOLDE^118^, and REFMAC5. Figures were generated using PyMOL and ChimeraX.

### In vitro GCase activity measurements

GCase specific activity was measured by incubating varying concentrations of purified recombinant GCase (imiglucerase, Sanofi) with a fixed concentration of 4-Methylumbelliferyl β-D-glucopyranoside (4-MUG, Thermo Fisher 10815-812). For specific activity measurements, 500mM stocks of 4-MUG (prepared in DMSO) were diluted to 20mM in assay buffer (100 mM phosphate citrate, 0.5% sodium taurocholate, 0.25% Triton X-100, pH 5.2). An 11-point titration of GCase, from 25 nM–0.024 nM, was made by 2-fold serial dilution of GCase with assay buffer, in a 96 well polypropylene V-bottom plate. Samples were transferred to a black 384-well Nunc plate (Thermo Fisher Cat #262260) in 5 uL duplicates and equal volume of 20 mM 4-MUG was added to all samples, which were sealed and incubated for 1 hour at room temperature. The reaction was quenched with the addition of 10 uL “stop buffer” (0.5 M sodium carbonate, 0.5 M sodium bicarbonate, pH 10.3), and the plate was read using a Biotek Synergy Neo2 multimode plate reader using excitation wavelength 365 nm and emission wavelength 445 nm. The fluorescence signals were interpolated against a 4-MU standard curve with a concentration range of 0 pmoles – 400 pmoles to determine the amount of product formed from the reaction.

To measure GCase enzyme kinetics, 2nM GCase was assayed against an 11-point 2-fold serial dilution of 4-MUG, ranging from 36 mM down to 0.035 mM, with a 12th well containing only assay buffer as the buffer blank. GCase enzyme samples were diluted to 4 nM in assay buffe, and the reaction was initiated by adding an equal volume of GCase to the 4-MUG titration at 15 minutes, 30 minutes, 45 minutes, 60 minutes, or 90 minutes. All reactions were stopped at the same time via addition of the stop buffer in equal volume to the reaction volume, and plates were read using a Biotek Synergy Neo2 multimode plate reader using excitation wavelength 365 nm and emission wavelength 445 nm.

### Cell free liposome assay

The neutral liposome composition consisted of 5% cholesterol, 10% PE, 75% PC, and 10% GlcCer and the 30% BMP liposome composition contained 30% BMP, 5% cholesterol, 10% PE, 45% PC and 10% GlcCer by weight. The starting lipids were purchased from Avanti Polar Lipids and were solubilized in chloroform and normalized to 10 mg/mL, after which the lipid solutions were combined in their respective proportions according to volume. The chloroform was then evaporated from the solution using nitrogen gas; solutions were rotated in a round bottom flask to ensure that the lipids dried in an even thin film and were further dried under vacuum overnight. Lipids were resuspended by addition of 1x TBS to normalize the lipid concentrations to 20 mg/mL and water bath sonication (approximately 5 minutes). Liposome solutions were then aliquoted into Eppendorf tubes and stored at -20°C. Assays were performed in a buffer consisting of 100 mM sodium acetate pH 5.3 and 150 mM NaCl, unless otherwise stated. Liposome-substrate working solutions were prepared by sonication of liposome stocks, in a water bath, until the stock solutions were optically clear, followed by addition of assay buffer. The GCase reactions were initiated by the addition of executed commercially available Cerezyme (imiglucerase, Sanofi) or recombinant GCase proteins to the liposome substrate solution in an AlphaPlate 384-shallow well (Revvity; #6008350). The final working concentrations used in the assay were 50 nM GCase and 100 µM GlcCer. The primary reaction was incubated at 37°C for 1 hour at 180 rpm. The secondary reaction started by addition of Glucose-Glo reagent (Promega, #J6022) which was prepared according to manufacturer instructions and added to the reaction wells. Addition of the Glo reagent also resulted in the quenching of the primary reaction. The secondary reaction was left to incubate at 37°C for 1 hour at 180 rpm. The luminescence was quantified using a PerkinElmer Envision plate reader.

### Sample lysis & immunoblotting

HEK cell lysates were prepared in RIPA buffer supplemented with SDS to a final concentration of 2%, cOmplete protease inhibitor (Roche, 04693159001), phosSTOP (Roche, 04906837001), and Benzonase nuclease (Sigma-Aldrich, E1014). Human iPSC-derived cell lysates and mouse brain lysates were prepared in Cell Lysis Buffer (Cell Signaling Technology, #9803), supplemented with cOmplete protease inhibitor, phosSTOP, and Benzonase nuclease. All samples were prepared for SDS-PAGE analysis by addition of NuPAGE LDS Sample Buffer (Thermo, NP0007) and NuPAGE Sample Reducing Agent (Thermo, NP0004). For all blots, samples were first resolved on a 4-12% Bis-Tris NuPAGE gel (Thermo) in 1x MOPS running buffer (Thermo, NP0001) and then transferred to nitrocellulose membranes (Bio-Rad) using the Trans-Blot Turbo Transfer System (Bio-Rad). Membranes were blocked for at least 1h at room temperature in Intercept (TBS) Blocking Buffer (LI-COR, 927-60001) and then incubated in primary antibody solution (prepared in Intercept Blocking Buffer + 0.05% Tween-20) overnight at 4°C. Membranes were rinsed in TBS-T, incubated in secondary antibody solution (prepared in Intercept Blocking Buffer + 0.05% Tween-20) for 1h at room temperature, and then rinsed extensively with TBS-T before being imaged on and Odyssey CLx Infrared Imaging System (LI-COR).

### LysoIP

Lysosomes were isolated from HEK cells stably over-expressing the TMEM192-HA^x3^ tag as previously described^119^ with the following modifications. Cells were harvested and resuspended in KPBS+ buffer and then fractionated by passing the suspension through a 23g needle five total times. Intact nuclei and other heavy membranes were pelleted by centrifugation (800g, 10min), and lysosomes were immunoprecipitated from the resulting post-nuclear supernatant (PNS) by incubation with anti-HA magnetic beads (Thermo, 88837) with end-over-end rotation for 15 minutes. Lysosome-bound beads were washed 2x with KPBS+ buffer and 2x with KPBS buffer (3-minute incubation with end-over-end rotation in each wash). Lysosomes were eluted from beads for immunoblotting by resuspending beads in NuPAGE LDS sample buffer + NuPAGE Sample Reducing Agent, followed by incubation at 95°C for 10 minutes. Lysosomes were eluted from beads for proteomic analysis by incubation in PBS supplemented with 5% SDS, with 2000rpm shaking for 15 minutes at room temperature. Lysosomal lipids were eluted from the beads for lipidomic analysis as described below.

### EndoH Experiment

Glycan removal by EndoH was performed according to guidelines from manufacturer (New England Biolabs (NEB), P0702). Unless otherwise noted, all steps were performed on ice or at 4°C using pre-chilled buffers and reagents. Cell lysates from HEK cells were prepared in Cell Lysis Buffer (Cell Signaling Technology, #9803), supplemented with cOmplete protease inhibitor, phosSTOP, and Benzonase nuclease, and insoluble material was cleared by centrifugation (21,000g, 10min). Total protein concentration of the cleared lysates was measured via BCA assay (Thermo, 23225), and lysates were normalized to the same total protein concentration using lysis buffer. Lysates were then denatured by mixing 99ul of lysate with 11ul of 10x Glycoprotein Denaturing Buffer (NEB), followed by incubation at 100°C for 10 minutes. 25ul of denatured lysates was mixed with 4ul of 10x GlycoBuffer 3 (NEB) and 2ul of EndoH (NEB) in a final reaction volume of 40ul, followed by incubation at 37°C for 6h. Treated lysates were prepared for SDS-PAGE/immunoblotting and glycosylation content of endogenous GCase present in the lysates was assessed by mobility shifts in resulting immunoblots.

### Strep-Tactin pulldown

Synthesis and cloning into the pCDNA3.1(+) vector (Thermo V79020) of GBA1 cDNA sequences (either WT or E326K) with a c-terminal StrepII tag was performed at Genscript. GBA KO HEK cells, plated in 15cm dishes the day prior, were transfected with 20µg GBA1-WT-StrepII or 25µg GBA1-E326K-StrepII plasmids using Lipofectamine 3000 (Thermo, L3000001) in normal growth media according to the manufacturer’s protocol. Growth media was completely replaced the following day, and cells were further incubated for a total of 48h post-transfection before being assayed. Cells were harvested and resuspended with ice-cold 1x PBS to rinse, and the resulting cell suspension was split into two equal portions before being pelleted by centrifugation. Cell pellets were lysed in lysis buffer base comprised of 0.5% NP-40, 150mM NaCl, 5mM EDTA, cOmplete protease inhibitor (Roche), phosSTOP (Roche), and pH was adjusted by addition of either 50mM HEPES (pH 7.3) or 50mM MES (pH 5.5), for 30min at 4°C with end-over-end rotation. Lysates were cleared of insoluble material by centrifugation (21,000g, 10min, 4°C), and 60µl of MagStrep Strep-Tactin XT beads (IBA Lifesciences, 2-5090-002, pre-washed in corresponding pH lysis buffer) were added to equal volumes of the clarified lysates. Beads were incubated with lysates at 4°C with end-over-end rotation for 2 hours to ensure complete binding, followed by 3x washes (5 minutes each, at 4°C with end-over-end rotation) in lysis buffer (of corresponding pH). Bound proteins were eluted from beads by resuspending beads in NuPAGE LDS sample buffer + NuPAGE Sample Reducing Agent, followed by incubation at 95°C for 10 minutes. Samples were analyzed by immunoblotting, and LIMP2 band intensity in pulldown samples was expressed relative to the total GCase (or StrepII) band intensity within each sample, and values from each sample in an experimental replicate were normalized to the WT-GBA1-transfected sample lysed at pH 7.3 from that same replicate.

### Lipidomics Analysis

Cell lysate and lysosome samples were prepared in methanol:water (90:10, v/v) extraction buffer containing stable-isotope and surrogate internal standards. Samples were shaken for 20 minutes at room temperature and then incubated for 1 hour at -20°C. Samples were centrifuged at 18,000 x g for 5 minutes at 4°C. For lipidomics analysis, 50% volume of the resulting supernatant was analyzed by LC-MS. For glycosphingolipid analysis, 25% volume of the resulting supernatant was dried under N₂ gas, then reconstituted in acetonitrile:isopropanol:water (92.5:5:2.5, v/v/v) fortified with 5 mM ammonium formate and 0.5% formic acid for LC-MS analysis. Lipid and glycosphingolipid analyses were conducted using liquid chromatography (Agilent Infinity II 1290) coupled with electrospray mass spectrometry (QTRAP 6500+). Lipids were analyzed in both positive and negative modes on a UPLC BEH C18 column (150 × 2.1 mm, 1.7 μm, Waters Corp.) at 55 °C, with a 10-minute gradient at 0.25 mL/min, as previously described.^55^ Glycosphingolipids were analyzed in positive mode on a HALO HILIC column (150 × 3.0 mm, 2.0 μm, Advanced Materials Technology) at 45 °C, with an 8.5-minute gradient at 0.48 mL/min, also as previously described.^55^ Quantification and peak integration were performed using MultiQuant 3.3 (ABSciex) with a minimum signal-to-noise (S/N) ratio of >5 and points across baseline >8. Analyte abundances were calculated as area ratios of endogenous lipids to class-level stable isotope standards and exported for statistical analysis.

### Proteomics Analysis

Cell lysate samples were prepared in PBS with 5% SDS. IP samples were eluted off HA magnetic beads after enrichment with PBS with 5% SDS. Both samples were processed using S-trap (Protifi) 96-well plate format according to manufacturer’s protocol. Briefly, samples were reduced with the addition of 15 mM TCEP and incubated on a thermomixer at 60°C for 30 mins and 1000 rpm. Samples were then cooled to room temperature, IAA was added to 15 mM and incubated in the dark for 1 hr. Samples were then acidified to 1.2% phosphoric acid and 7 volumes of S-trap binding buffer (50 mM TEAB, 90% methanol pH 7.2) was added. Samples were loaded on the S-trap plate at 1500xg for 2 min, then washed 3x with 200 uL S-trap binding buffer. Trypsin was added at 1:30 in 100 mM TEAB and incubated on a thermomixer at 37°C overnight. Samples were eluted stepwise with TEAB, 0.2% formic acid, then 0.2% formic acid/50% acetonitrile. The eluates were combined and dried in a speedvac. Dried peptide was resuspended in 0.1% formic acid, measured by nanodrop and adjusted to 0.5 ug/uL. Samples were analyzed using an Agilent 1290 fitted with split-flow nanoflow adapter coupled to a Bruker timsTOF Pro 2. Peptide (0.5 ug) was loaded on an Aurora Ultimate column 25cm x 75um ID with 1.7um (IonOpticks) with the flowrate set to approximately 300 nl/min at 98% mobile phase A (0.1% formic acid in water) and 2% mobile phase B (0.1% formic acid in acetonitrile) with a gradient of 22% B at 90 mins, 37% B at 105 min, 80% B from 115-120 min then reconditioning at 2% B between 120-145 min. The MS was set to 100 ms ramp time with a duty cycle of 100%, dia-PASEF settings were set to a mass range of 300-1400 Da with an Ion mobility range of 0.70-1.40 1/K0 and collision energy scaling from 20eV at 0.60 1/K0 to 65 eV at 1.60 1/K0. DIA windowing was determined based on prior runs using py_diAID.^120^ Raw data was analyzed using Spectronaut version 19 (Biognosys). Raw files were converted into the Spectronaut format and calibrated using a global spectral library, with processed data including detailed peptide and protein quantification metrics for downstream analysis.

### Lipidomics & proteomics data analysis

Lipid and protein features detected in at least 70% of samples were retained. Partially missing lipid and protein values were imputed using the kNN function from the VIM package or minProb function from the MsCoreUils package, respectively. For lipidomics, unwanted technical variations unrelated to biological factors were removed using the RUV4 function from the ruv package. Differentially regulated lipids and proteins were analyzed with a robust linear model, incorporating experimental groups (e.g., genotype) and covariates (e.g., batch). Group-wise contrasts were evaluated using lmFit and contrast.fit, followed by robust Empirical Bayes moderation (logFC ≥ 1.1, FDR via Benjamini-Hochberg). Visualizations (box plots, volcano plots, heatmaps) were generated using ggplot2, EnhancedVolcano, and pheatmap. Feature overlap and Venn diagrams were analyzed with VennDiagram, while GSEA was conducted using clusterProfiler (gene sets: 3–800, FDR adjusted via Benjamini-Yekutieli). All analyses were performed in R (4.2.1).

### Seahorse mitochondrial respiration assay

Three independent clones of WT, GBA1-p.E326K, GBA1-p.L444P and GBA1 KO HEK293 cells (15,000 cells/well) were seeded on PDL-coated 96-well Agilent Seahorse XFe96/XF Pro PDL Cell Culture microplate (Cat #103799-100) in DMEM (Thermo Cat # 11965092) + 10% FBS media for 24 h. For the mitochondrial respiration experiments, the Seahorse XF Cell Mito Stress Test Kit (Cat # 103015-100) was used. One day before Seahorse experiment, Extracellular Flux Assay cartridge (Cat #103792-100) was hydrated with 200 ul/well DI water and placed in a non-CO2 incubator at 37 C. On the day of the assay, cells were washed twice with assay media comprised of XF DMEM (Cat # 103575-100), 10 mM glucose (Cat # 103577-100), 1 mM pyruvate (Cat # 103578-100) and 2 mM Glutamine (Cat # 103579-100). Water in Extracellular Flux Assay cartridge was also replaced with pre-warmed 200 ul/well Seahorse XF calibrant (Cat # 100840-000) and placed back in non-CO2 incubator at this time. Cells were imaged using brightfield microscopy to obtain cell counts utilized for normalization. Cells were then incubated for 1 hour in a non-CO2 incubator before beginning the Seahorse experiment. Ports on the sensor plate were filled according to the XF Cell Mito Stress Test Kit and cells were subjected to sequential injections of oligomycin (final concentration 2 uM), FCCP (1 uM), and rotenone/antimycin A (0.5 uM each). Data was analyzed using the Agilent Seahorse Analytics online software to generate kinetic curves and calculate maximal respiration.

### TMRM Staining

HEK cells were seeded in the inner 60 wells of PDL-coated 96-well imaging plates the day prior to staining in normal culture media. Cells were stained for 30 minutes at 37°C in culture media supplemented with 100nM TMRM (Thermo I34361), 100nM MitoTracker Green (Thermo M7514), and 2µM Hoechst 33342. Cells were washed into imaging buffer comprised of 1x DPBS supplemented with 20mM HEPES, 5% FBS, and 5.55mM D-glucose, and imaged on an automated confocal high-content imager (Revvity, Opera Phenix Plus High-Content Screening System) using a 63x water immersion objective lens, 405nm, 488nm, and 561nM excitation lasers, and preset emission filters. Channel acquisition was separated to avoid fluorescence spillover between channels. A custom analysis in the Harmony 5.2 software (Revvity) was used to quantify TMRM and MitoTracker Green signal from the resulting images. Cell nuclei were segmented and counted from Hoechst stain using the “Find Nuclei” building block, cell boundaries were segmented from MitoTracker Green signal using the “Find Cytoplasm” building block, and the sum cytoplasmic TMRM and MitoTracker Green signal was calculated using the “Calculate Intensity Properties” building block, before being normalized to the number of nuclei. Mitochondrial membrane potential was expressed as the ratio of normalized TMRM/MitoTracker Green signal, measured from 12 non-overlapping images per well. A total of four individual wells per cell line were imaged per plate, and the average mitochondrial membrane potential across all four wells is reported for a single experimental replicate.

### Animal Care & Sourcing

For animals used for brain lipidomics analysis, all animal procedures were performed in accordance with a protocol approved by the Institutional Animal Care and Use Committee of the National Institute on Aging, NIH. Wildtype, homozygous and heterozygous E326K GBA1 knock-in (KI) male and female mice raised on a C57Bl/6 background were bred in-house on a 12-hr day/night cycle for the following experiments. All mice were supplied with Rodent NIH-07 diet and water ad libitum.

For animals used for analysis of GCase activity & enzyme levels, all procedures were performed with adherence to ethical regulations and protocols approved by Denali Therapeutics Institutional Animal Care and Use Committee. Mice were housed under a 12-hour light/dark cycle and had access to water and standard rodent diet (#25502, irradiated; LabDiet) ad libitum. Gba^E326K^ mice (and age-matched WT control mice) were obtained from the Jackson Laboratory (strain #036159).

### Preparation of mouse brain lysates for lipidomics

Approximately 20 mg of flash-frozen brain tissue was homogenized in 20 volumes of methanol extraction buffer containing stable-isotope and surrogate internal standards. Homogenization was performed using a 3 mm tungsten carbide bead with a Qiagen TissueLyzer II for 30 seconds at 25 Hz, repeated twice. The lysate was centrifuged at 21,000 x g for 20 minutes at 4°C. The supernatant was collected and stored at -20°C for 1hr to allow for further precipitation, followed by a second centrifugation at 4,000 g for 20 minutes at 4°C. Aliquots for both lipidomics and glycosphingolipid analyses were recovered, processed, and analyzed as described above for cell lysate samples.

### Measurement of GCase activity from mouse brain lysate

Brain tissue (∼50 mg) was homogenized in 10X volume (ul) by tissue weight (mg) cold lysis buffer (1ml of 10X cell lysis buffer (9803S Cell Signaling Technology), 9 ml of distilled water, 1 tablet cOmplete protease inhibitor (Roche 04693116001), and 1 tablet PhosSTOP protease inhibitor (Roche 04906845001)) with a 3-mm stainless steel bead using the Qiagen TissueLyzer II for two rounds of 3 min at 27 Hz. Homogenates were then centrifuged at 17,000 rpm for 15 min at 4°C and the supernatant collected. The resulting lysate was stored at −80°C.

Lysates were thawed on ice, mixed by inversion and centrifuged briefly. Lysates were then pre-diluted by adding 10ul to 150ul of assay buffer (100 mM Phosphate Citrate Buffer, pH 5.2) and mixed by gentle pipetting. 4ul of diluted lysate was then added to 86ul of assay buffer containing 1mM 4-MUG substrate (Sigma M3633-1G), mixed at 700 rpm for 5 min on an orbital shaker and transferred to a 37°C incubator for 1hr. 100ul of stop solution (500 mM Glycine, 300 mM NaOH, pH 9.8) was then added to the reaction and fluorescence was read on a BioTek plate reader.

### LysoFQ-GBA flow cytometry assay in dissociated mouse brains

Mice were anesthetized with Avertin and perfused with ice cold PBS for 10min. Brains were collected and placed into ice cold DPBS (Gibco 14040133) with 0.5% BSA (Miltenyi 130-093-376). On parafilm on ice, cerebellum and olfactory bulb were removed and brain tissue was minced into 3mm chunks. Tissues were dissociated using the Adult Brain Dissociation Kit (Miltenyi Biotec, 130-107-677) following manufacturer’s protocol. Dissociated cells were kept at 4°C during all steps unless noted. The resulting cell pellet was resuspended in 400ul of assay buffer (0.5%BSA in DPBS), and 20ul of cell suspension was taken for microglia counting in a 1.5ml Protein Lobind tube by staining with 80ul of staining solution (FACS Buffer PBS (Gibco 10010023) + 1% BSA (Miltenyi 130-093-376), CD11b - BV421 1:100 (BD Biosciences 562605), CD45 - APC 1:100 (BD Biosciences 559864), FC block 1:100 (Biolegend 101320)). Cells were incubated on ice under foil for 15min and 500ul of FACS buffer was added. Counting samples were centrifuged 300g 5min 4C and the supernatant aspirated. Pellets were resuspended in 50ul of FACS buffer with Propidium Iodide (Miltenyi 130-093-233) (1:200) and added to the bottom of the FACS tube without filtration. 50ul of CountBright Absolute Counting Beads (Invitrogen, C36950) was added and microglia concentration determined using the BD ARIA III flow cytometer.

Microglia concentrations were normalized with assay buffer and 50,000 microglia in 100ul per sample were added to individual reaction tubes (1.5ml protein lobind). 100ul of 22uM LysoFQ-GBA probe in assay buffer was then added to each tube. Reactions were mixed by pipetting gently and placed in a 37°C incubator for 1hr. Tubes were placed back on ice and 1ml of assay buffer added to wash followed by a 5min centrifugation at 300g. The supernatant was aspirated, and pellets resuspended in 100ul of staining solution (FACS Buffer PBS (Gibco) + 1% BSA (Miltenyi 130-093-376), CD11b - BV421 1:100 (BD Biosciences 562605), CD45 – APC 1:100 (BD Biosciences 559864), FC block 1:100 (Biolegend 101320) and incubated 25min covered with foil. 1ml of FACS buffer was added to each tube and centrifuged 300g 5min. The supernatant was aspirated, and pellets resuspended in 300ul FACS buffer with PI (Miltenyi 130-093-233) (1:200). Samples were loaded into FACS tubes through the 30um filter and probe fluorescence was measured with FACS ARIA III recording 3000 live microglia per sample. Data analysis was performed in FlowJo (BD Biosciences) and mean probe fluorescence values for microglia were obtained.

### iPSC line generation & microglial differentiation

GBA1 KO and knock-in of the E326K mutation to the GBA1 locus was performed by Thermo Fisher Scientific Life Technologies Custom Services. Parental iPS cells were transfected (via electroporation) with TrueCut Cas9 V2 (Thermo A36498), gRNA, and ss-Oligo donor template to induce desired edits. Clonal cell populations were isolated and homozygous knock-in or knock out was confirmed by Sanger sequencing of the target locus. For GBA1 KO the gRNA sequence 5’-GCGGTGTGTCTCCCCTAGGG-3’ and ss-Oligo sequence 5’-AOZCCCTCCCTCCCAGGTGCCCGCCCCTGCATCCCTTAAAGCTTCGGCTACAGCTCG GTGGTGTGTGTCTGCAATGCCACATACTGTGACTCCTTTGFOC-3’ was used, and for E326K knock-in the gRNA sequence 5’-ACATGGTACAGGAGGTTCTA-3’ and ss-Oligo sequence 5’-AOZTGGAGCCCACACAGGCCTCTGAGGCAAAGAGCATGGTGTTGGGGAACAGGCGG TGTGTCTTCCCTAGGGTGGCTTTGGCTGGAGCCAGAAAGTCOFG-3’ was used. Pluripotency of resulting clones was determined using the TaqMan hPSC Scorecard (Thermo A16179), and clones were confirmed to have same genetic background via karyotyping using the KaryoStat+ with Cell ID assays (Thermo A52986).

Hematopoietic progenitor cells (HPCs), were generated using an adapted protocol.^121^ iPSCs from the lines generated above were maintained on matrigel coated plates in mTeSR+ (Stemcell technologies #100-0276) at no greater than 80% confluency prior to starting differentiation. On Day 1 cells were lifted with Accutase (Stemcell Technologies #07930) and seeded onto Vitronectin (Thermo Fisher, A31804) coated plates at 5.5-6e^5^ cells/cm^2^. Cells were seeded in E8 (Stemcell Technologies #05990) containing Activin A 10ng/mL (Peprotech #120-14P), BMP4 40ng/mL (Peprotech #120-05), CHIR99021 3µM (Tocris #4423/10) and Y-27632 10µM (ATCC #ACS-3030). On Day 2, approximately 18hrs later, Day 1 medium was aspirated, and E6 (Stemcell Technologies # 05946) containing Activin A 10ng/mL, BMP4 40ng/mL and IWP2 2µM (Selleckchem #S7085) was added. On Day 3, approximately 24hrs later, Day 2 medium was aspirated, and E6 containing Activin A 10ng/mL, BMP4 40ng/mL, IWP2 2µM and thermostable FGF2 20ng/mL (Millipore Sigma #GF446) was added. On Day 4 medium was aspirated and cells were accutase treated for approximately seven minutes to singularize them. Cells were pelleted and a small sample was stained using anti-CD235a antibody 1:100 (BD #551336) following standard flow cytometry analysis practices. If the cell sample was greater than 20% positive for CD235a expression the remaining cells were replated onto Matrigel-coated plates into STEMdiff Hematopoietic Medium B (Stemcell Technologies #05310) containing Y-27632 10µM (ATCC #ACS-3030). A complete medium change was performed. From this point forward the vendor protocol was followed until Day 12 cell collection. On Day 12 cells were collected from suspension, pelleted and cryopreserved as batches of HPCs. A fraction of cells was retained for flow cytometry analysis. If cells were less than 95% positive for anti-CD43 (Biolegend #343204) those lots were not used for microglial differentiation.

All microglial differentiation was performed from cryopreserved stocks of HPCs. Prior to thawing HPC vials plates were coated with matrigel and STEMdiff Microglia Differentiation Kit medium was prepared (Stemcell Technologies #100-0019). Cells were thawed using standard practices and cryoprotectant was washed away using DMEM/F-12 (ThermoFisher #11320033) and by pelleting cells at 500xg for five minutes. HPCs were resuspended in microglial differentiation medium and plated at a density of 1.5-2e^5^ cells/cm^2^. Maximum density did not exceed 2e^6^ cells/60cm^2^. From this point forward the vendor protocol was followed with only minor adjustments to accommodate schedules where some of the medium additions were omitted. Additionally, no replating was performed until Day 24 when cells were transitioned to STEMdiff Microglia Maturation Medium (Stemcell Technologies #100-0020). At this stage fresh matrigel coated plates were used and cells were transitioned and maintained according to the vendor’s protocol. Mature iMicroglia were assessed for anti-CD45 (BD #561863) and anti-CD11b (BD #557321) purity, by flow cytometry analysis, prior to use for all experiments with a target percentage of 90% double positive.

### Human genetics analysis of whole blood GCase activity and plasma metabolite levels

We used publicly available data from PPMI to examine the relationship of GBA1 p.E326K and GBA1 PD pathogenic variants to GCase activity and GCase substrates, relative to non-carriers. Whole blood-derived GCase activity was available via PPMI Project 132. We restricted the analysis to samples from the baseline visit (N = 439). We next sought to further restrict this cohort to samples from subjects of predicted European ancestry. To do this, we utilized whole genome sequencing data available through the Accelerating Medicines Partnership – Parkinson’s Disease (AMP-PD) data resource (2021_v2-5release_0510; N = 10,418) and restricted the data to subjects from PPMI (N = 1,807). We then merged this data with 1000 Genomes Phase 3 (1000G) reference samples (PMID: 26432245) and performed linkage disequilibrium (LD)-based pruning of this merged dataset followed by principal component analysis. These analyses were carried out using plink v1.9 (PMID: 25722852). We then used the first 10 principal components and 1000G super population labels to predict genetic ancestry for all samples in the PPMI dataset via the k-nearest neighbors algorithm. A total of 394 of the 439 samples were predicted as European ancestry and retained for downstream analysis. To classify subjects based on p.E326K and GBA1 pathogenic variant status, we used the consensus *GBA1* and *LRRK2* coding variant summary also available through PPMI. 19 subjects were classified as having p.E326K, 14 subjects as having a pathogenic variant (including 8 p.N370S carriers, 2 p.L444P carriers, and 1 carrier each of IVS2+1G>A and p.R463C, and 1 compound heterozygote of p.T369M and p.R120W and 1 homozygous N370S carrier), and 361 having neither. In the latter group, 3 subjects were missing PD case/control status, 1 subject was missing *GBA1* variant status, and 11 subjects had *GBA1* variants that were not considered as pathogenic in the consensus coding document (p.R39C and p.T369M) and were excluded from the analysis, leaving 346 subjects in this group.

Measures for 330 plasma metabolites were available via PPMI Projects 180 and 215 on a total of 640 unique subjects. Measures from the first available visit were used in downstream analyses. 584 subjects were of predicted European ancestry (as described above) and retained for further analysis. Of these, 14 were determined to be p.E326K carriers (extracted from the WGS data), 159 were classified as having a *GBA1* pathogenic variant (including 105 p.N370S carriers), 410 were classified as having neither, and 1 was missing from the PPMI consensus coding summary and removed from further analysis.

For both whole blood GCase activity and all plasma metabolite measurements, separate models were constructed to compare p.E326K carriers to non-carriers, and GBA1 pathogenic variant carriers to non-carriers. The outcome variables were natural log-transformed and fit in a linear model consisting of age, sex, disease status (PD vs control), and the first 3 principal components derived from genome-wide genotype date (post LD-based pruning, again using plink 1.9 (PMID: 25722852)) to account for potential confounding by ancestry. The residuals from these models were then inverse normal transformed using the blom() function from the “rcompanion” R package and used in a linear model to evaluate the association between p.E326K or GBA pathogenic variant status vs. non-carriers.

## Data Availability

Whole blood GCase activity data is available through PPMI (Project ID: #132). Plasma metabolite data is also available through PPMI Project ID: #180 and GCase activity and progranulin measurements through PPMI Project # 215. WGS data from PPMI is available through AMP-PD (https://www.amp-pd.org/register-for-amp-pd). Coordinates and cryoEM maps for the structure of the GCase E326K-LIMP2 complex have been deposited in the PDB and EMDB with the following accession codes: PDB 9P70 (model), EMD-71330 (maps). Coordinates and related data for the crystal structure of GCase E326K have been deposited in the PDB with the accession code 9P71.

## Acknowledgements

Data used in the preparation of this article were obtained from the Accelerating Medicine Partnership® (AMP®) Parkinson’s Disease (AMP PD) Knowledge Platform. For up-to-date information on the study, visit https://www.amp-pd.org. The AMP® PD program is a public-private partnership managed by the Foundation for the National Institutes of Health and funded by the National Institute of Neurological Disorders and Stroke (NINDS) in partnership with the Aligning Science Across Parkinson’s (ASAP) initiative; Celgene Corporation, a subsidiary of Bristol-Myers Squibb Company; GlaxoSmithKline plc (GSK); The Michael J. Fox Foundation for Parkinson’s Research; Pfizer Inc.; Sanofi US Services Inc.; and Verily Life Sciences. ACCELERATING MEDICINES PARTNERSHIP and AMP are registered service marks of the U.S. Department of Health and Human Services. Parkinson’s Progression Markers Initiative (PPMI)– a public-private partnership – is funded by the Michael J. Fox Foundation for Parkinson’s Research and funding partners, including 4D Pharma, Abbvie, AcureX, Allergan, Amathus Therapeutics, Aligning Science Across Parkinson’s, AskBio, Avid Radiopharmaceuticals, BIAL, BioArctic, Biogen, Biohaven, BioLegend, BlueRock Therapeutics, Bristol-Myers Squibb, Calico Labs, Capsida Biotherapeutics, Celgene, Cerevel Therapeutics, Coave Therapeutics, DaCapo Brainscience, Denali, Edmond J. Safra Foundation, Eli Lilly, Gain Therapeutics, GE HealthCare, Genentech, GSK, Golub Capital, Handl Therapeutics, Insitro, Jazz Pharmaceuticals, Johnson & Johnson Innovative Medicine, Lundbeck, Merck, Meso Scale Discovery, Mission Therapeutics, Neurocrine Biosciences, Neuron23, Neuropore, Pfizer, Piramal, Prevail Therapeutics, Roche, Sanofi, Servier, Sun Pharma Advanced Research Company, Takeda, Teva, UCB, Vanqua Bio, Verily, Voyager Therapeutics, the Weston Family Foundation and Yumanity Therapeutics. The PPMI investigators have not participated in reviewing the data analysis or content of the manuscript. Data used in the preparation of this article was obtained on October 9^th^, 2023 from the PPMI database (www.ppmi-info.org/access-dataspecimens/download-data), RRID:SCR_006431. For up-to-date information on the study, visit www.ppmi-info.org. The investigators within the LCC contributed to the design and implementation of the LCC and/or provided data and/or collected biospecimens, but did not necessarily participate in the analysis or writing of this report. We thank Jamal Alkabsh for support for LC-MS/MS based analyses and members of the lysosomal function pathway team at Denali for useful discussions and feedback. We thank Marcus Chin for useful feedback on image analysis. We thank Joseph Lewcock, Chris Koth, Thomas Sandmann and Karen Lai for useful feedback on the manuscript. The synchrotron data was collected at the P13 beamline operated by EMBL Hamburg at the PETRA III storage ring (DESY, Hamburg, Germany).

## Authors’ contributions

O.B.D., L.R., J.E.K., S.V.A., N.S.G., and J.H.S. contributed to the conception, design, acquisition, analysis, data interpretation of this work and contributed to manuscript writing. S.S.D., R.G., J.H.K., E.T., J.P.C., M.T.M., N.E.P., and J.C.U. contributed to the conception, design, acquisition, analysis and data interpretation of this work. H.P.B. contributed to the analysis and interpretation of the data. M.A., A.H.N., H.N.N., E.I.L., T.Y., K.X., R.A.P.S., R.L., S.B., contributed to the acquisition, analysis, and data interpretation of this work. M.P., A.N.Q., X.W., S.Z., G.D.P., M.S.K., C.S.M., A.A., D.J.V., and M.R.C., contributed to the conception and design of the work. A.G.H. contributed to the conception, design, interpretation of data, and manuscript writing.

