## Supplementary Materials for "A Common PD-Risk *GBA1* Variant Disrupts LIMP2 Interaction, Impairs Glucocerebrosidase Function, and Drives Lysosomal and Mitochondrial Dysfunction"

**Affiliations:**

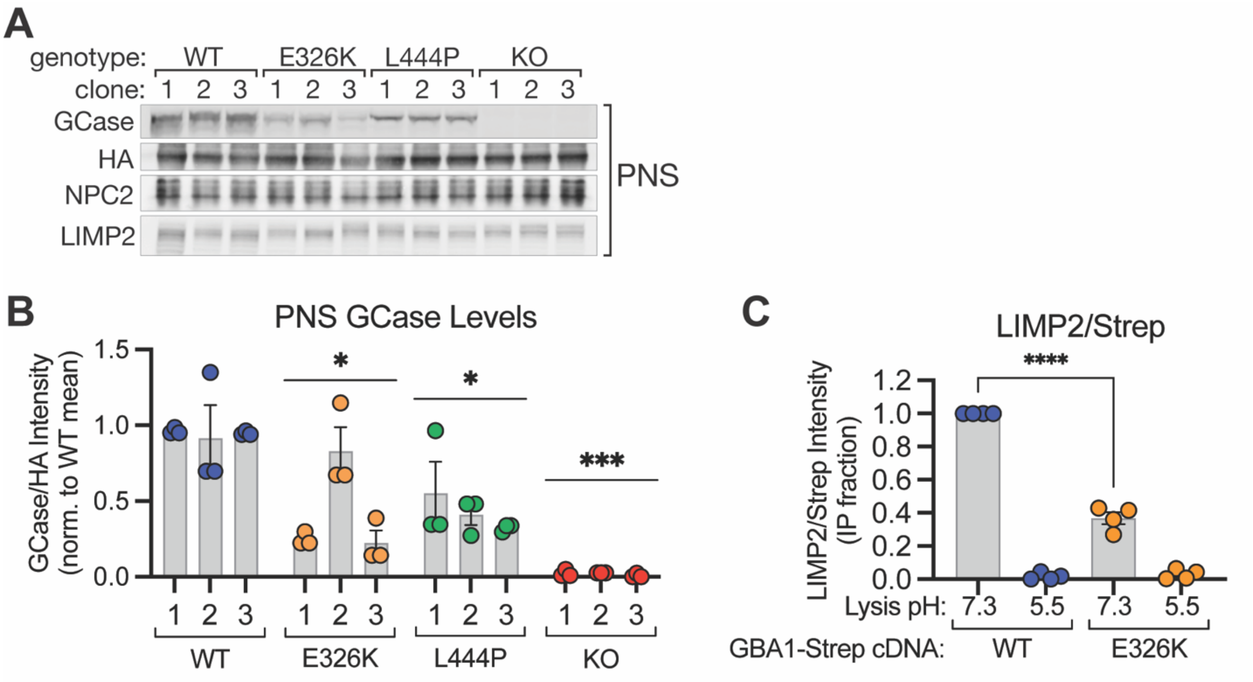

**Figure S1. Related to data in Figure 1. A** Representative immunoblot of post-nuclear supernatant (PNS) samples corresponding to Lyso-IP samples shown in Fig. 1E. **B** Quantification of GCase levels in PNS samples (normalized to HA band intensity), corresponding to Lyso-IP quantification shown in Fig. 1F. n = 3 replicates. **C** Quantification of LIMP2:Strep band intensity ratios, normalized to WT-GBA1/pH 7.3 lysis condition within each replicate, from pulldown fraction of experiment shown in Fig 1I. n = 4 replicates. All bar graphs shown mean ± SEM with individual points representative of data from a single experimental replicate. Unless otherwise noted, all statistics were performed on genotype-level comparisons using one-way ANOVA with Dunnett's multiple comparison test, where * = $p < 0.0332$, *** = $p < 0.0002$, **** = $p < 0.0001$.

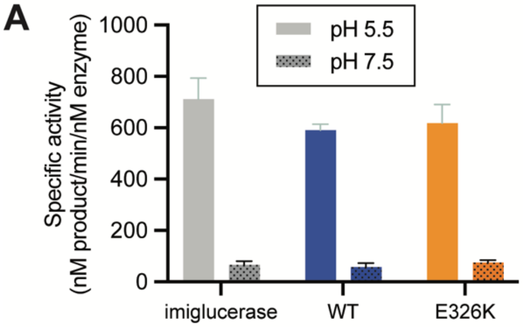

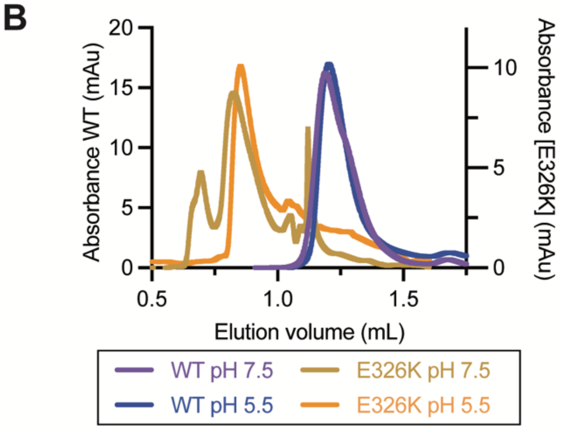

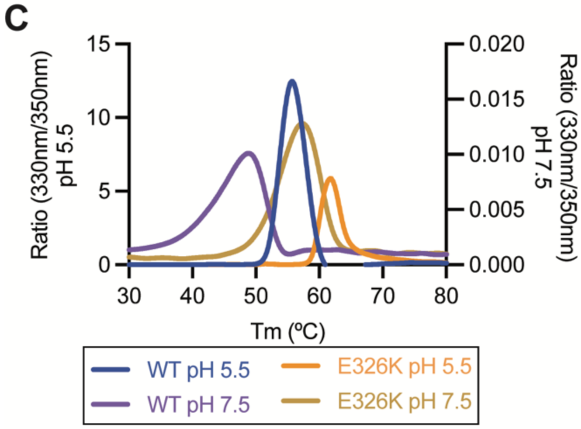

**D**

| | | pH 5.5 | pH 7.5 | Difference |
|---|---|---|---|---|
| $T_m$(ºC) | WT | 54.4 | 43.6 | 8.1 |
| | E326K | 60.1 | 54.5 | 5.6 |
| DLS radius (nm) | WT | 4.4 | 4.7 | -0.32 |
| | E326K | 8.6 | 7.1 | 1.57 |

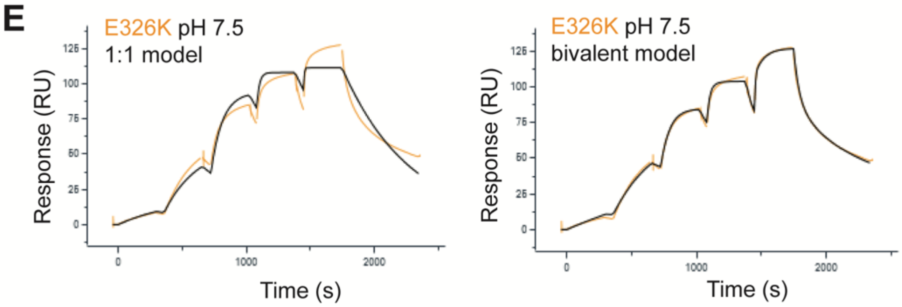

**F**

| biotinylated ligand | pH | immobilization level (RU) | theoretical Rmax (RU) | experimental Rmax (RU): WT GCase | experimental Rmax (RU): E326K GCase |
|---|---|---|---|---|---|
| LIMP2 | 5.5 | 96.5 | 110.3 | 99.8 | 164.0 |
| | 7.5 | 79.5 | 90.9 | 70.6 | 113.2 |

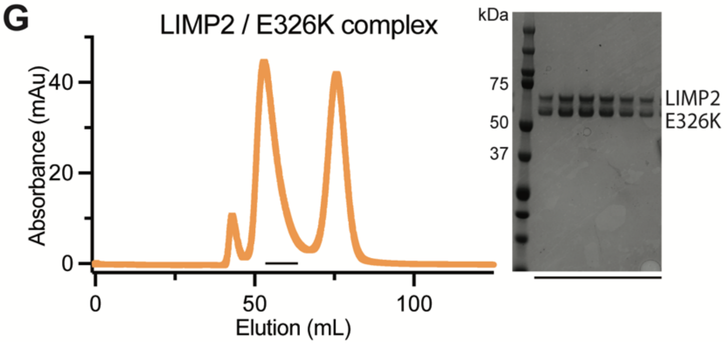

**Figure S2. Related to data in Figure 2. A** Activity against 4-MUG substrate of WT (blue) and E326K variant (orange) measured at neutral (stippled bars) and acidic (solid bars) pH and compared to imiglucerase (grey). Lack of activity at neutral pH is expected and consistent across all three constructs. **B** Overlay of size exclusion chromatography profiles for both WT GCase and E326K variant at pH 5.5 or pH 7.5. Delipidated samples were run on a S200 pre-equilibrated with 25mM HEPES, 150mM NaCl, pH 7.5 or 20mM Acetate, 150mM NaCl, pH 5.5. **C** Thermomelt profiles measured for WT GCase and E326K variant at neutral and acidic pH using a NanoDSF. **D** Summary table for biophysical characterization of recombinant WT GCase or E326K variant. Represented values correspond to the average of 3 separate duplicates for DLS (nm) and $T_m$ (ºC). **E** Representative SPR sensograms of E326K binding to biotinylated LIMP2 at neutral pH. Recombinant E326K was injected in increasing concentrations over surface immobilized LIMP2. Responses were fitted to a 1:1 (left) or bivalent (right) model. **F** Comparison between theoretical and experimental Rmax values; theoretical values were calculated based on molecular weight and immobilization level while Rmax values reflect average responses from two replicate injections. Higher Rmax values for E326K suggest altered binding stoichiometry or complex formation consistent with a multivalent interaction. **G** Size exclusion chromatography profile of GCase E326K complexed with an excess of LIMP2. Fractions containing the complex were run on SDS-PAGE to assess purity.

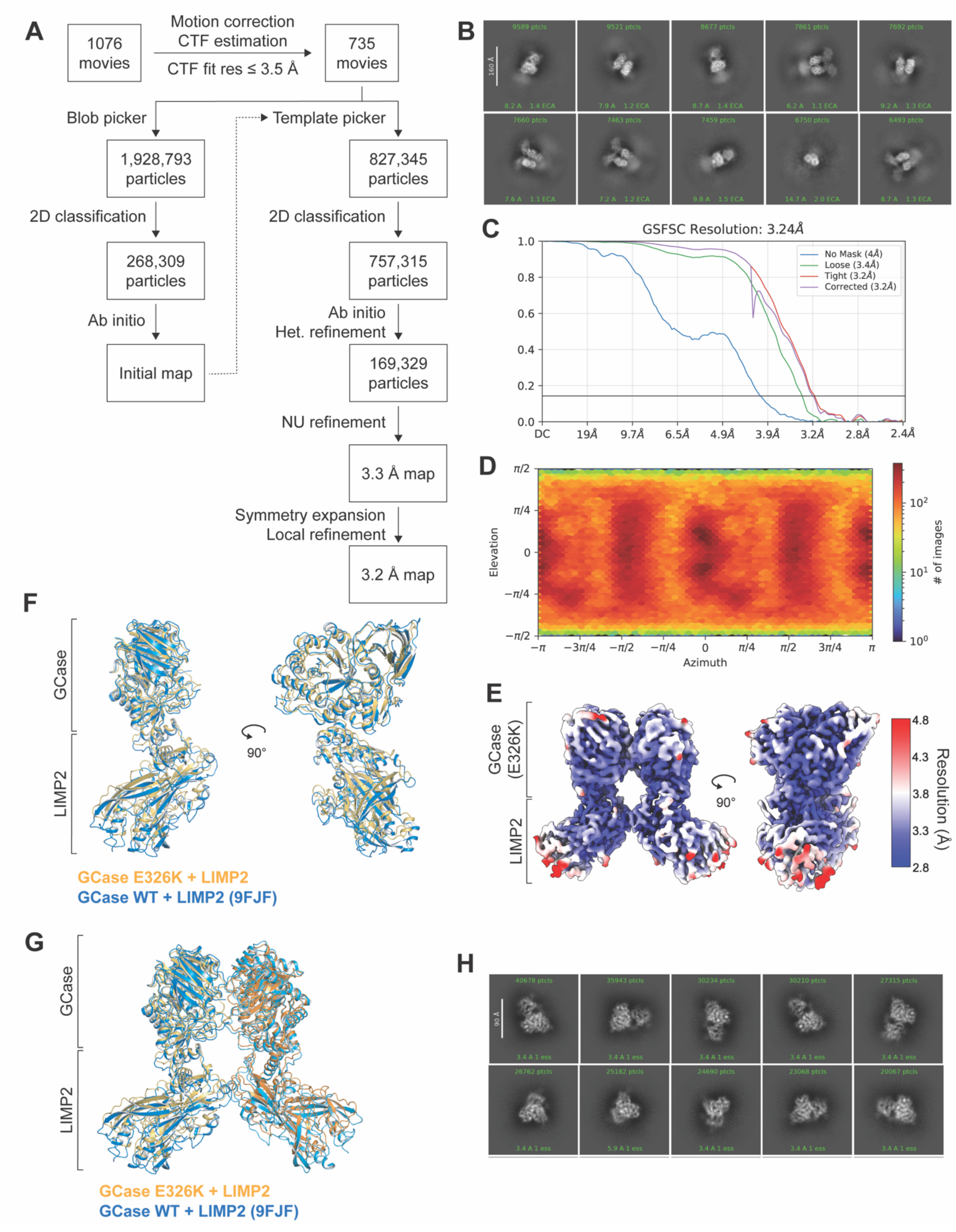

**Figure S3. Related to data in Figure 2. A** Single-particle image processing workflow for GCase E326K/LIMP2 reconstruction. **B** Representative 2D classes averages of GCase

E326K/LIMP2 dataset. **C** Fourier Shell Correlation (FSC) curve for final refinement. **D** Heat map representation of particle orientation distribution. **E** CryoEM map of GCase E326K/LIMP2 complex colored by local resolution. **F** Overlay between GCase E326K/LIMP2 complex and published WT GCase/LIMP2 cryoEM structure (PDB: 9FJF, nanobodies used as fiducial markers were removed for clarity). The complexes were aligned on the GCase portions of the structures **G** Alignment of two copies of GCase WT/LIMP2 heterodimer from 9FJF with each protomer in the GCase E326K/LIMP2 structure. The complexes were aligned on the GCase portions of the structures **H** Representative 2D classes averages of GCase WT/LIMP2 dataset.

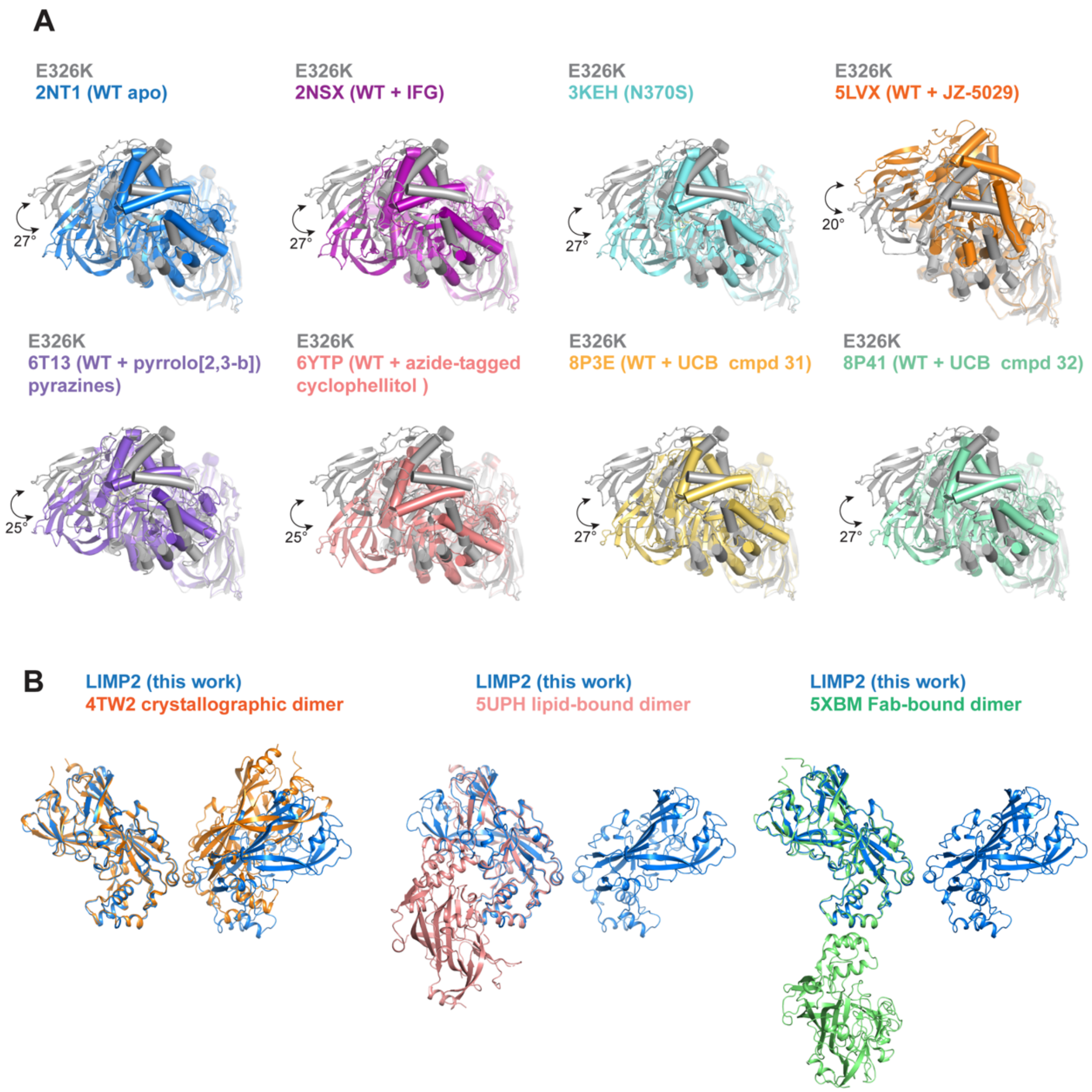

**Figure S4. Related to data in Figure 3. A** Overlay between observed E326K dimer (grey) in our E326K-LIMP2 cryoEM structure and published crystallographic dimers including; WT apo GCase (PDB:2NT1), WT GCase bound to pharmacological chaperone isofagomine (PDB:2NSX), Gaucher variant N370S (PDB:3KEH) and several compound-bound structures. Figures were generated by aligning one protomer in each crystallographic dimer to chain A of the E326K dimer, shown in the back of each image, to demonstrate the offset of the other protomer with chain B of the E326K dimer, shown in the front. Alignments show that the dimers are slightly rotated relative to each other and the difference in angle between the unaligned protomers in the front are annotated. The difference in angle was calculated in PyMOL using the angle between domains script (https://raw.githubusercontent.com/speleo3/pymol-psico/master/psico/orientation.py) **B** Overlay between observed LIMP2 homodimer from GCase E326K-LIMP2 cryo-EM structure (blue) and crystallographic LIMP2 dimers from the following PDBs: 4TW2, 5UPH and 5XBM. Dimers were obtained by generating symmetry related molecules from published structures.

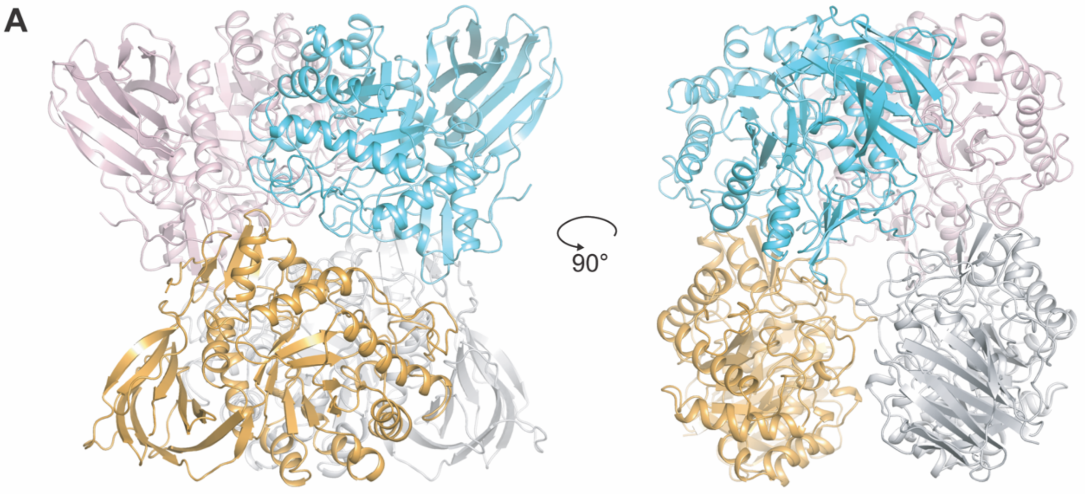

**B**

| | DLS (nm) | | | $T_m$ (ºC) | |
|---|---|---|---|---|---|
| | radius pH 5.5 | radius pH 7.5 | difference | pH 5.5 | pH 7.5 |
| WT | 4.4 | 4.7 | -0.32 | 54.4 | 46.3 |
| E326K | 8.6 | 7.1 | 1.57 | 60.1 | 54.4 |
| E326K+R329D dimer | 9.3 | 6.6 | 2.7 | 57.0 | 49.1 |
| E326K+R329D monomer | 7.6 | 6.7 | 0.83 | 54.1 | 46.1 |
| E326K+R329E dimer | 9.9 | 6.5 | 3.4 | 57.4 | 48.6 |
| E326K+R329E monomer | 6.2 | 6.1 | 0.04 | 54.4 | 46.2 |
| E326K+R329A dimer | 10.7 | 9.3 | 1.42 | 57.1 | 50.7 |

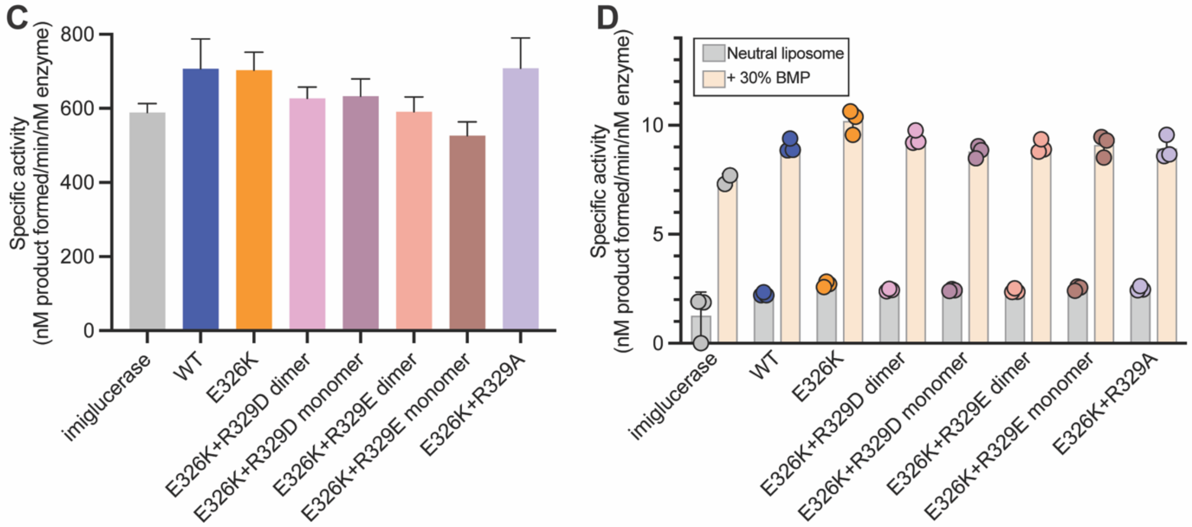

**Figure S5. Related to data in figure 3. A** Cartoon representation of the organization of GCase molecules within the asymmetric unit. Molecules are organized as a dimer of dimers. Physiologically relevant dimers are colored in grey/orange and cyan/pink. **B** Summary table comparing protein radii and melting temperature of double mutants in comparison with WT and E326K constructs at neutral and acidic pH. Monomeric and dimeric species were isolated post SEC purification and tested individually. **C** Activity of double mutants compared to imiglucerase (gray), WT (blue), and E326K variant (orange) measured at acidic pH in 4-MUG assay. **D** Enzymatic activity of purified recombinant

GCase variants against GlcCer in liposomes prepared with either neutral lipids (gray bars) or with 30% BMP (orange bars).

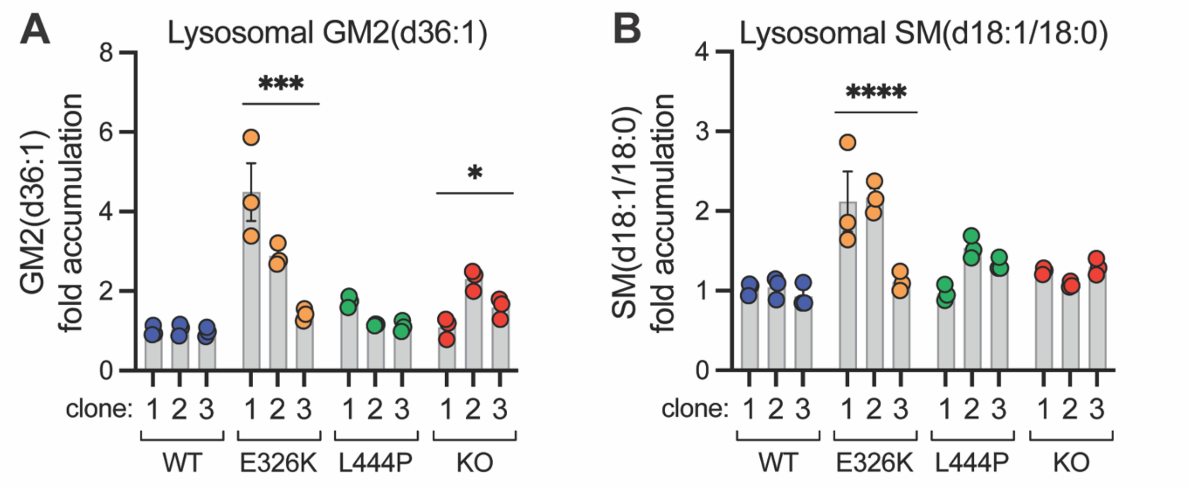

**Figure S6. Related to data in Figure 4. A** Fold change (over WT mean) of GM2(d36:1) in Lyso-IP samples from HEK293T cells of the indicated genotype, related to data shown in Fig. 4D. n = 3 replicates. **B** Fold change (over WT mean) of SM(d18:1/18:0) in Lyso-IP samples from HEK293T cells of the indicated genotype, related to data shown in Fig. 4D. n = 3 replicates. All bar graphs shown mean ± SEM with individual points representative of data from a single experimental replicate. Unless otherwise noted, all statistics were performed on genotype-level comparisons using one-way ANOVA with Dunnett's multiple comparison test, where * = $p < 0.0332$, *** = $p < 0.0002$, **** = $p < 0.0001$.

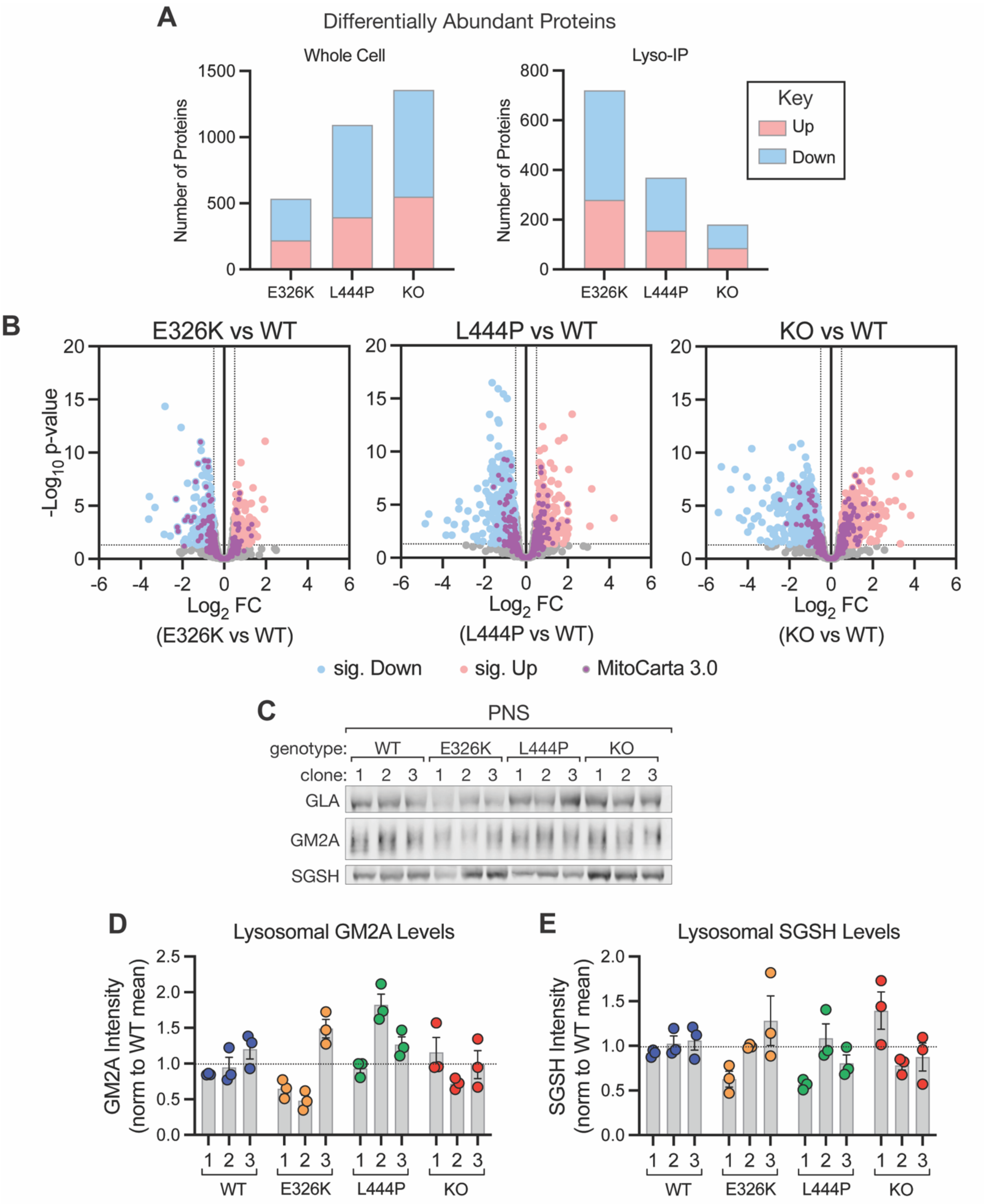

**Figure S7. Related to data in Figure 5. A** Number of differentially abundant proteins ($p < 0.05$; Absolute $Log_2FC \geq 0.5$; relative to WT cells or lysosomes) from proteomic analysis of whole cell or Lyso-IP samples from HEK293T cells of the indicated genotype. Red bars indicate number of proteins with increased abundance, and blue bars indicate number of proteins with decreased abundance, relative to WT. Related to data shown in Fig. 5A, B. n = 3 replicates. **B** Volcano plots showing differential abundance of proteins identified in whole cell samples from indicated GBA1 variant cells compared to WT cells. Proteins with significantly changed abundance ($p < 0.05$; Absolute $Log_2FC \geq 0.5$) are highlighted in red or blue for increased or decreased abundance, respectively. Proteins identified as mitochondrial in origin in the MitoCarta 3.0 database are highlighted in purple. Related to data shown in Fig 5A, B. n = 3 replicates. **C** Representative immunoblot of PNS samples corresponding to Lyso-IP samples shown in Fig. 5G. **D** Quantification of GM2A levels (normalized to HA band intensity) measured via immunoblotting of Lyso-IP fraction & normalized to mean of all WT samples. Corresponding to data shown in Fig. 5G. n = 3 replicates. **E** Quantification of SGSH levels (normalized to HA band intensity) measured via immunoblotting of Lyso-IP fraction & normalized to mean of all WT samples. Corresponding to data shown in Fig. 5G. n = 3 replicates. For A & B, statistical tests were performed on genotype-level comparisons using robust linear model as described in methods and significance threshold was set at FDR adjusted p value < 0.05. Bar graphs in D-E show mean ± SEM with individual points representative of data from a single experimental replicate.

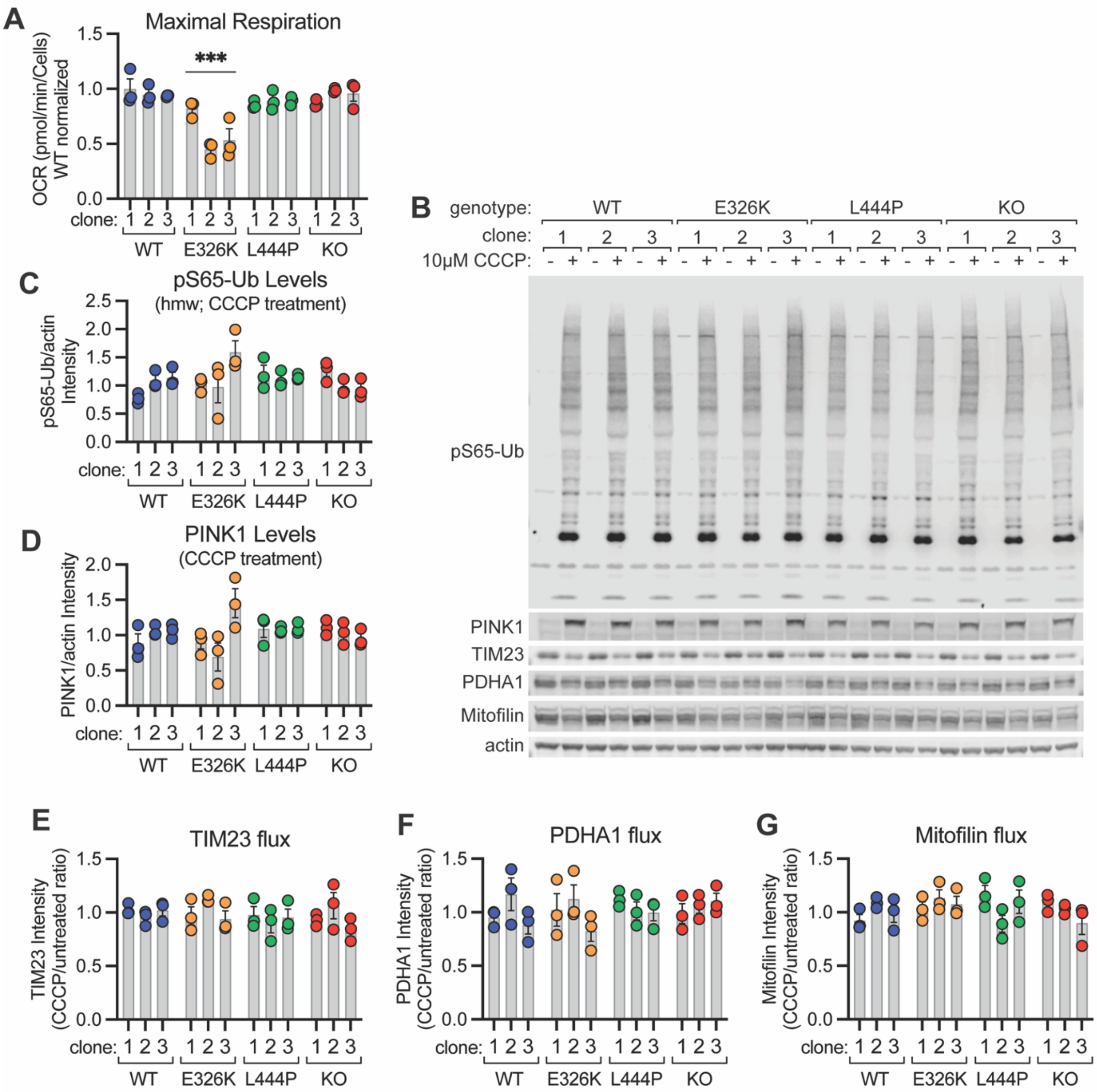

**Figure S8. Related to data in Figure 5. A** Maximal respiratory capacity (quantified by OCR) of HEK293T cells of indicated genotype, measured using Seahorse XF Cell Mito Stress Test assay. Data corresponds to OCR traces shown in Fig 5D. n = 3 replicates. **B** Representative immunoblot of HEK293T cells of indicated genotype treated with 10µM CCCP for 16 hours. **C, D** Quantification of high molecular weight (hmw) pS65-Ub (C) or PINK1 (D) band intensity, normalized to actin band intensity, from CCCP-treated samples, measured via immunoblot & normalized to mean of all WT cells **E-G** Quantification of CCCP-induced turnover (“flux”; the ratio of CCCP-treated:untreated band intensity) of TIM23 (E), PDHA1 (F), or Mitofilin (G), measured via immunoblot & normalized to the mean flux of all WT cells. For C-G, n = 3 replicates. All bar graphs show mean ± SEM with individual points representative of data from a single experimental replicate. Unless otherwise noted, all statistics were performed on genotype-level comparisons using one-way ANOVA with Dunnett’s multiple comparison test, where *** = $p < 0.0002$.

### Table S1. Cryo-EM data collection, refinement and validation statistics

| | GCase E326K-LIMP2 (EMD-71330) (PDB 9P70) |
|---|---|
| **Data collection and processing** | |
| Magnification | 120,000x |
| Voltage (kV) | 200 |
| Electron exposure (e–/Å$^2$) | 50 |
| Defocus range (μm) | -2.0 to -1.0 |
| Pixel size (Å) | 1.2 |
| Symmetry imposed | C2 |
| Initial particle images (no.) | 1121820 |
| Final particle images (no.) | 153182 |
| Map resolution (Å) | 3.2 |
| FSC threshold | 0.143 |
| Map resolution range (Å) | 2.8 - 47 |
| **Refinement** | |
| Initial model used (PDB code) | 2NT1, 4TW2 |
| Model resolution (Å) | 3.2 |
| FSC threshold | 0.5 |
| Model resolution range (Å) | 2.8 - 47 |
| Map sharpening *B* factor (Å$^2$) | -96.5 |
| Model composition | |
| Non-hydrogen atoms | 14464 |
| Protein residues | 1792 |
| Ligands | 0 |
| *B* factors (Å$^2$) | |
| Protein | 88.9 |
| R.m.s. deviations | |
| Bond lengths (Å) | 0.005 |
| Bond angles (°) | 1.740 |
| Validation | |
| MolProbity score | 1.67 |
| Clashscore | 1.75 |
| Poor rotamers (%) | 2.33 |
| Ramachandran plot | |
| Favored (%) | 92.72 |
| Allowed (%) | 6.94 |
| Disallowed (%) | 0.34 |

**Table S2. Data collection and refinement statistics (molecular replacement)**

| PDB ID | GCase E326K<br>9P71 |
|---|---|
| **Data collection** | |
| Space group | P 43 21 21 |
| Cell dimensions | |
| *a*, *b*, *c* (Å) | 132.1 132.1 340.7 |
| α, β, γ (°) | 90.00, 90.00, 90.00 |
| Resolution (Å) | 55.11 – 3.07 (3.148 – 3.068) |
| $R_{sym}$ or $R_{merge}$ | 0.210 (10.073) |
| $I / \sigma I$ | 6.9 (0.8) |
| CC1/2 (%) | 0.987 (0.292) |
| Completeness (%) | 97.79 (86.5) |
| Redundancy | 11.4 (12.0) |
| **Refinement** | |
| Resolution (Å) | 3.07 |
| No. reflections | 53351 |
| $R_{work}$ / $R_{free}$ | 0.248/0.283 |
| No. atoms | |
| Protein | 15874 |
| Ligand/ion | 296 |
| Water | 2 |
| *B*-factors | |
| Protein | 86.13 |
| Ligand/ion | 103.17 |
| Water | 66.79 |
| R.m.s. deviations | |
| Bond lengths (Å) | 0.0092 |
| Bond angles (°) | 1.484 |

*Values in parentheses are for highest-resolution shell.

**Table S3. Full statistics for comparison of GBA1 p.E326K carriers relative to non-carriers for association with plasma metabolite and protein measurements in PPMI (Project #215).** Analytes were log transformed, fit in a linear model consisting of age, sex, disease status (PD vs control), and the first 3 principal components derived from genome-wide genotype data from the samples specific to each analysis. Residuals were then inverse-normal transformed and used in a linear model against carrier status (see Methods). Estimate = standard deviation change in adjusted and transformed analyte levels in carriers relative to non-carriers. SE = standard error. Stat = test statistic. P = p-value. L95, U95: 95% confidence interval. N = number of samples that went into analysis. Values less than sample counts reported in the methods reflect missingness in analyte levels.

**Table S4. Full statistics for comparison of GBA1 pathogenic variant carriers relative to non-carriers for association with plasma metabolite and protein measurements in PPMI (Project #215).** Analytes were log transformed, fit in a linear model consisting of age, sex, disease status (PD vs control), and the first 3 principal components derived from genome-wide genotype data from the samples specific to each analysis. Residuals were then inverse-normal transformed and used in a linear model against carrier status (see Methods). Estimate = standard deviation change in adjusted and transformed analyte levels in carriers relative to non-carriers. SE = standard error. Stat = test statistic. P = p-value. L95, U95: 95% confidence interval. N = number of samples that went into analysis. Values less than sample counts reported in the methods reflect missingness in analyte levels.
